# Hibiscus bullseyes reveal mechanisms controlling petal pattern proportions that influence plant-pollinator interactions

**DOI:** 10.1101/2024.02.05.579006

**Authors:** Lucie Riglet, Argyris Zardilis, Alice L. Fairnie, May T. Yeo, Henrik Jönsson, Edwige Moyroud

## Abstract

Colourful patterns on flower corollas are key signals to attract pollinators. The formation of such motifs relies on the establishment of developmental boundaries that partition the growing petal epidermis into different subdomains, where cells can produce specific pigments and acquire distinctive cell shapes and textures. While some of the transcription factors and biosynthetic pathways producing these characteristics as cell differentiate have been extensively studied, the upstream processes restricting the activities of molecular players to specific regions of the petal epidermis remain enigmatic. Here, we unveil that the petal surface of *Hibiscus trionum*, an emerging model system featuring a bullseye on its corolla, is pre-patterned as the position of the bullseye boundary is specified long before the motif becomes visible to the human eye. Using a 1-D computational model, we explore how a boundary established at such an early stage can be maintained throughout development. Reciprocally, by exploiting transgenic lines and natural variants, we show that plants can regulate the relative position of the boundary during the pre-patterning phase or modulate division and growth on either side of this boundary at later developmental stages to yield variations in final bullseye proportions. Finally, we provide evidence that such modifications in bullseye size have functional significance as buff-tailed bumblebees (*Bombus terrestris*) can reliably identify a food source based on the size of its bullseye. Notably, we found that individuals exhibit a clear preference for the larger bullseye of *H. trionum* over the smaller pattern of its close relative, *H. richardsonii*.

## Introduction

The petal epidermis of flowering plants showcases remarkable pattern diversity intricately tied to specialised functions. By combining regions that display contrasting features, such as colour, texture or cell morphology, these motifs play a crucial role in pollinator attraction, thus favouring plant reproduction and participating in speciation ^1–3^. Recent studies have also uncovered petal patterns that fulfil abiotic functions. UV-absorbing flavonoids can modulate transpiration, contributing to heat-retention and drought tolerance ^4–7^. This could explain why North American populations of common sunflowers (*Helianthus annuus*) found in colder, drier habitats tend to exhibit a larger UV-absorbing bullseye than those growing in warmer and more humid environments ^8^. Similarly, the size of the UV-absorbing centre on the corollas of distinct silverweed (*Argentina anserina)* populations correlates with the level of UVB irradiance they experience. Populations closer to the equator tend to display larger UV-absorbing bullseye, compared to those found at higher latitudes, possibly providing enhanced protection for pollen grains against UV damages ^9^. These size variations can evolve over extended time frames but also in response to environmental fluctuations driven by human activities^10^. Hence petal patterns likely represent dual adaptations to both biological and climatic factors but despite their functional significance the underlying processes governing the formation of these patterns remain poorly understood.

In developmental biology, a fundamental question remains: how are spatial patterns of distinct cell types specified and coordinated as tissues grow, ultimately giving rise to functional organs? Several elegant studies have investigated the regulatory processes that spatially control pigment production across the petal epidermis ^11–16^. Those repetitively singled out MYB and bHLH transcription factors, whose expression patterns account for the accumulation of pigments to specific portions of the petal epidermis. In contrast, our understanding of the formation and regulation of distinct cell shapes or cuticle textures across the petal surface remains limited ^17^. Achieving these characteristics seems to rely on precise spatio-temporal control of regulator expression (i.e., contribution of the MYB family to epidermis cell differentiation; Brockington et al., 2013) or different biosynthetic pathways (i.e., those involved in production of distinct cuticular components; ^18^. While these findings are valuable starting points, the upstream processes behind the restricted expression of genes orchestrating differentiation in neighbouring cells are still largely unexplored.

Here, we used the flower of *Hibiscus trionum* to explore the mechanistic basis of pattern formation. Its petals display a striking bullseye on the adaxial epidermis, with a purple-to-burgundy basal portion contrasting with a white surround (Fig. 1a). Further differences are found at the microscopic level: proximal epidermis cells, producing dark anthocyanin pigments, are flat, elongated, and covered with a striated cuticle, creating an iridescent blue-UV signal visible to pollinators ^19–21^. In contrast, distal cells are white, conical and their cuticle is smooth. Both regions are separated by a clear boundary, invariably located one-third from the petal base in wild-type (WT) individuals. How such a robust boundary is specified and then maintained while the petal is growing is not yet understood.

**Fig. 1.**
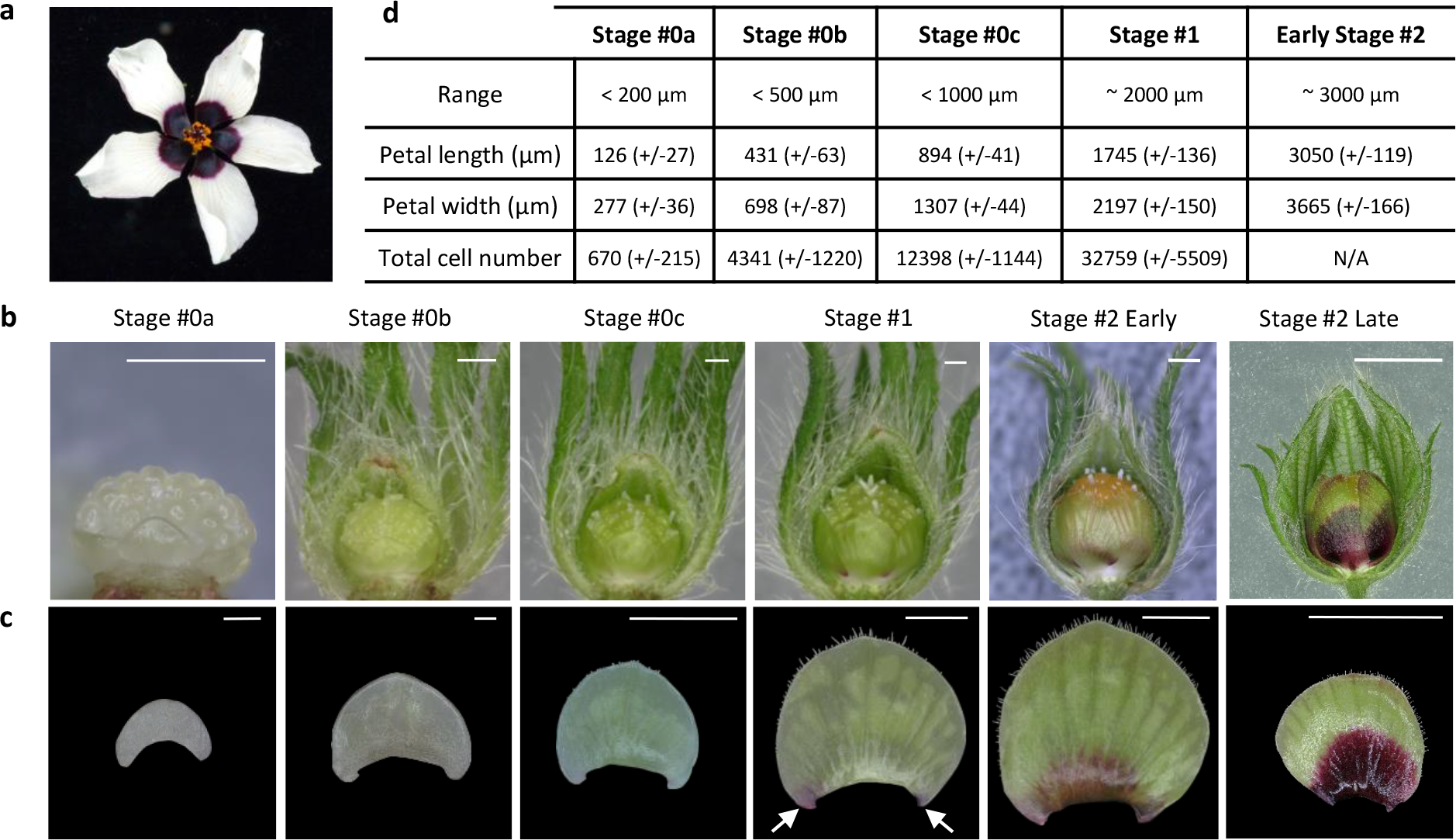
Early *Hibiscus trionum* petal development. **a**, Mature *H. trionum* flower (Stage 5). **b**, Petal organisation on flower buds ranging from Stage 0a (S0a) to Late Stage 2 (S2L). The images capture the abaxial side of the petal. Scale bars, 1 mm. **c**, Early developmental stages of the adaxial petal epidermis, from S0a to Early Stage 2 (S2E). Pigmentation emerges on both sides of the petal primordium at Stage 1 (S1), indicated by arrows. Scale bars, S0a and S0b : 100 μm, S0c to S2E : 1 mm, S2L : 5 mm. **d**, Classification criteria for *H. trionum* petal primordia. The total cell count was not assessed at S2E, as only the central petal stripe was imaged at that stage. n=5 petals for each stage.

*H. trionum* belongs to the Trionum complex, a group of Hibiscus species broadly distributed across Australasia ^22^. Within this complex, different species or populations exhibit a wide range of bullseye variations ^22^. The mechanisms driving such evolutionary changes in bullseye appearance are unknown and how such diversity impacts pollinator behaviour remains to be investigated.

To address these questions, we developed a comprehensive imaging pipeline to capture petal morphogenesis and analyse cell behaviour across the entire adaxial epidermis of Hibiscus petal during its development. We showed that even before any bullseye feature becomes apparent to the human eye, the petal is pre-patterned with the future bullseye domains already exhibiting differences in cell expansion and proliferation. Using a 1D-computational model, we also uncovered some of the developmental processes used by plants to maintain boundary position through growth or instead, to modify bullseye dimensions. Finally, we characterised the foraging behaviour of buff-tailed bumblebees (*Bombus terrestris*) in response to different bullseye proportions.

## Results

### The adaxial epidermis of *H. trionum* petal is pre-patterned

To start understanding how robust bullseyes form on the petal surface of *H. trionum* (Fig. 1a), we imaged the adaxial epidermis at fixed time points matching floral developmental stages ^18^. The dark purple-to-burgundy pigmentation is the first visible element of the pattern to emerge (Fig. 1b, c). At stage 0 (S0), the petal surface is entirely green without any noticeable sign of cellular differentiation (Fig. 1b, c). Colouration developing on both sides of the petal primordium attachment point to the floral structure is characteristic of stage 1 (S1) (Fig. 1b, c). By the end of early stage 2 (S2E), the basal region of the primordia is fully pigmented, and a sharp boundary separates the proximal and distal domains ^18^ (Fig. 1b, c). To pinpoint when and how this bullseye boundary emerges, we gathered quantitative cellular growth data across the petal epidermis at high spatial and temporal resolution. Until flowers open, petals are enclosed within the buds, with the growing adaxial petal epidermis (bearing the bullseye) facing inward, making it technically challenging to access (Fig. 1b). To generate a reference image dataset describing epidermis development at cellular resolution, we used confocal microscopy to capture images of dissected petals stained with FM1-43 to label cell outlines. After image segmentation using MorphographX ^23^, we quantified global changes in cell number and primordia dimensions (Fig. 1d, 2a). This analysis led us to subdivide the S0 phase into three sub-stages, stage 0a (S0a), 0b (S0b) and 0c (S0c) (Fig. 2a).

**Fig. 2.**
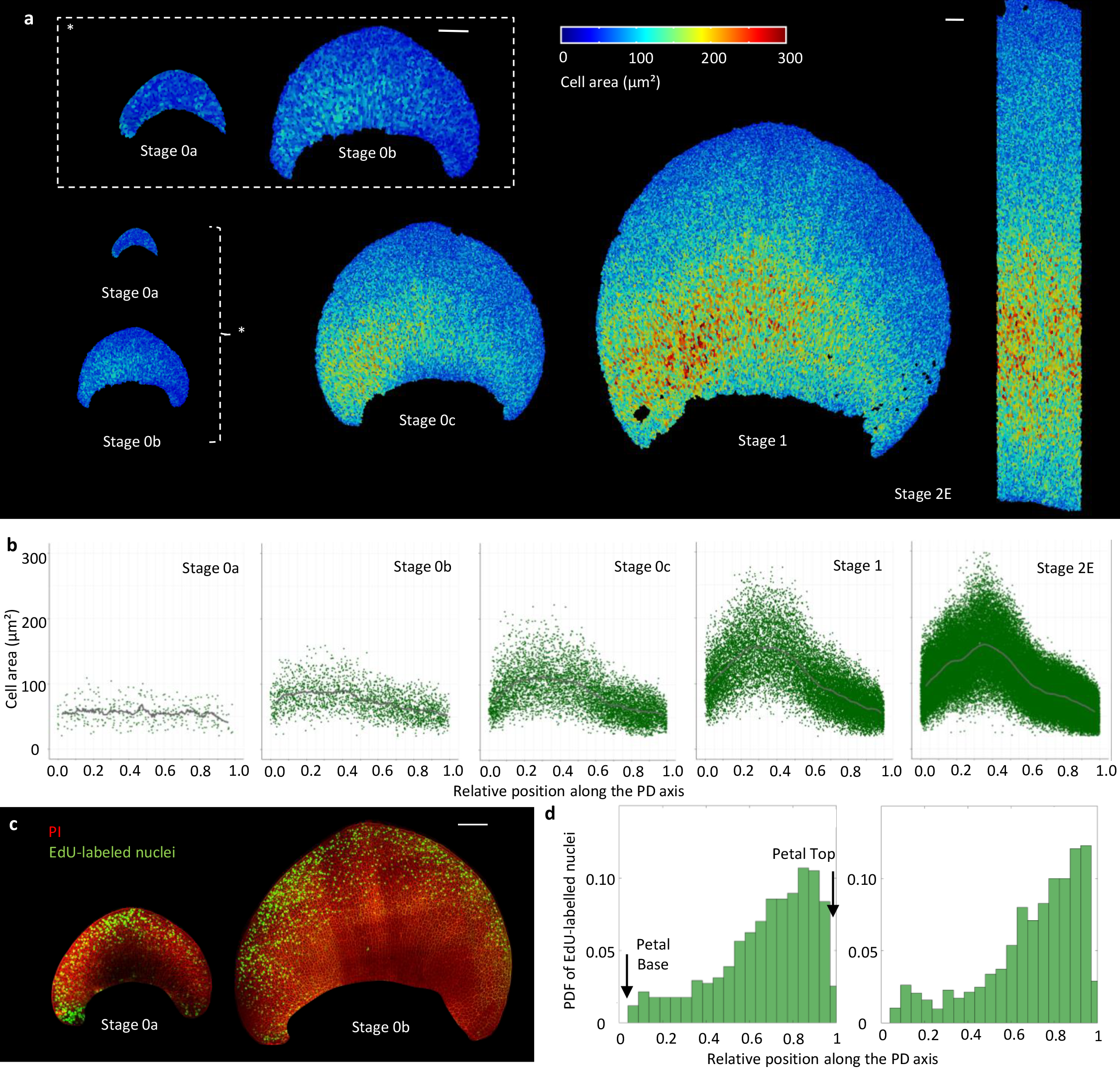
Spatio-temporal distribution of cell expansion and cell division events across the adaxial epidermis during the early stages of *Hibiscus trionum* petal morphogenesis. **a**, Colour map of cell area across the adaxial WT petal epidermis during early developmental stages (from S0a to S2E). Scale bar, 100 μm. **b**, Cell area distribution across the PD axis of *H. trionum* petal. The graphs consider only the central stripe of cells (20% of the petal width) for readability. Cell positions along the PD axis are relative, with 0 corresponding to the petal base, and 1 to the petal tip. Black lines correspond to the average cell area of all replicates. N = 5 petals for each stage. **c**, Distribution of cell division events across the adaxial epidermis of S0a and S0b petals. Newly synthesized DNA is labelled using fluorescently labeled nucleotide analog 5-ethynyl-2-deoxyuridine (EdU) (green) and plasma membranes are stained with propidium iodide (red). Scale bar, 100 μm. **d**, Probability density function (PDF) of the EdU-labeled nuclei along the PD axis of *H. trionum* S0a petals (stripes corresponding to 20% of the petal width and centered along the PD axis were analysed, see Supplement Fig. 1e). n = 5 petals for each stage.

We found that the cell area is uniform across the petal adaxial epidermis at S0a (Fig. 2a), but heterogeneity emerges at S0b as larger cells appear on one side of the petal. By S0c, the zone of larger cells expands from one petal side in a croissant-shaped pattern, resulting in right-left petal asymmetry (Fig. 2a). This motif becomes more pronounced towards S1 and S2E. The examination of cell area distribution along the proximo-distal (PD) axis, focusing on a central epidermis stripe (20% of the petal width), confirms that at S0a, cell area is, on average, uniform (Fig. 2b, Supplementary Fig. 1a). From S0b, cell areas are distributed heterogeneously, with larger cells preferentially located in the basal section of the adaxial petal epidermis. From S0c onward, cell area peaks around a third of the petal length from the base (Fig. 2b). This peak sharpens in S1, and the relative position of this cell size maximum along the PD axis is maintained as petal primordia grow to reach S2E (Fig. 2b). Additionally, from S1, cells in the proximal part of the petal epidermis elongate, already exhibiting a higher aspect ratio compared to those in the distal region (Supplementary Fig. 1b, c). Reciprocally, distal cells that will become conical at S4 ^18^ already display a higher circularity than proximal cells at S1 (Supplementary Fig. 1d). Thus, both cell area and geometry are regulated distinctly along the PD axis of the petal, with differences emerging along a croissant-shaped pattern during the S0 developmental phase, long before bullseye features (pigmentation, cuticular ridges and contrasting cell shapes) appear. Notably, this early pattern is characterised by a landmark at one third of the petal length, where the largest cells are located, which also matches with the position of the future bullseye boundary (S3 to S5; Supplementary Fig. 2a-e).

Next, we examined cell proliferation during the pre-patterning phase by mapping cell division events using a fluorescent nucleotide analog, 5-ethynyl-2-deoxyuridine (EdU) ^24^, which labels newly replicated DNA. We found that cell proliferation is not uniform across the S0a petal epidermis as most EdU-labelled nuclei reside in the distal region (Fig. 2c), mirroring the forthcoming bullseye layout. Quantifying the distribution of EdU-labelled nuclei along the PD axis of the petal (Fig. 2d,Supplementary Fig. 1e) confirmed that cell division events are mainly restricted to the upper half of the epidermis at S0a, with the highest proportion of fluorescent nuclei near the petal top (Fig. 2d). This distribution persists throughout the pre-patterning phase to S1 (Fig. 2c, d and Supplementary Fig. 1f, g). Thus, cell proliferation is differentially regulated across the two main regions of the bullseye (distal *vs*. proximal) very early during petal development.

Taken together, our results suggest that cell properties across the adaxial epidermis of the hibiscus petal are pre-patterned long before the bullseye distinctive features (pigmentation, cuticular ridges and contrasting cell shapes) become visible. This early pattern first emerges as a croissant-shaped distribution of cell size, with the largest cells laying at the one third mark along the petal PD axis and could already specify the position of the final bullseye boundary.

### The early pattern boundary coincides with the mature bullseye boundary

To understand the relationship between early pattern and bullseye formation, we analysed the distribution of cell features along the PD axis of the petal (Supplementary Fig. 2a-e) from late stage 2 (S2L) to stage 5 (S5, maturity) in *H. trionum* WT petals and tracked the position of the pigmentation boundary (corresponding to the transition from pink to white) (Supplementary Fig. 2d). The large size of these petals renders them unsuitable for cellular resolution imaging with our pipeline and segmentation using MorphographX. Instead, we manually measured cell features (area, aspect ratio, circularity) along a single line of cells from the base to top of the petal adjacent to the mid-vein and found that the pigmentation transition always matches changes in cell area (Supplementary Fig. 2e). We plotted the overall evolution of the boundary position (early pattern boundary form S0b to S2E and bullseye boundary from S2L to S5) across developmental stages in *H. trionum*. After its initial establishment one third from the petal base during the early patterning phase, the relative position of the boundary transiently rises to reach 0.4 at S2L before stabilising again around the one third mark at S4 (Fig. 3a). Hence, while proximal and distal regions exhibit very different growth properties, both in terms of cell division and expansion, the relative position of the boundary along the PD axis remains mostly constant throughout petal morphogenesis. This implies that growth differences between these regions are reconciled to maintain the relative lengths of the two zones that make the bullseye motif.

**Fig. 3.**
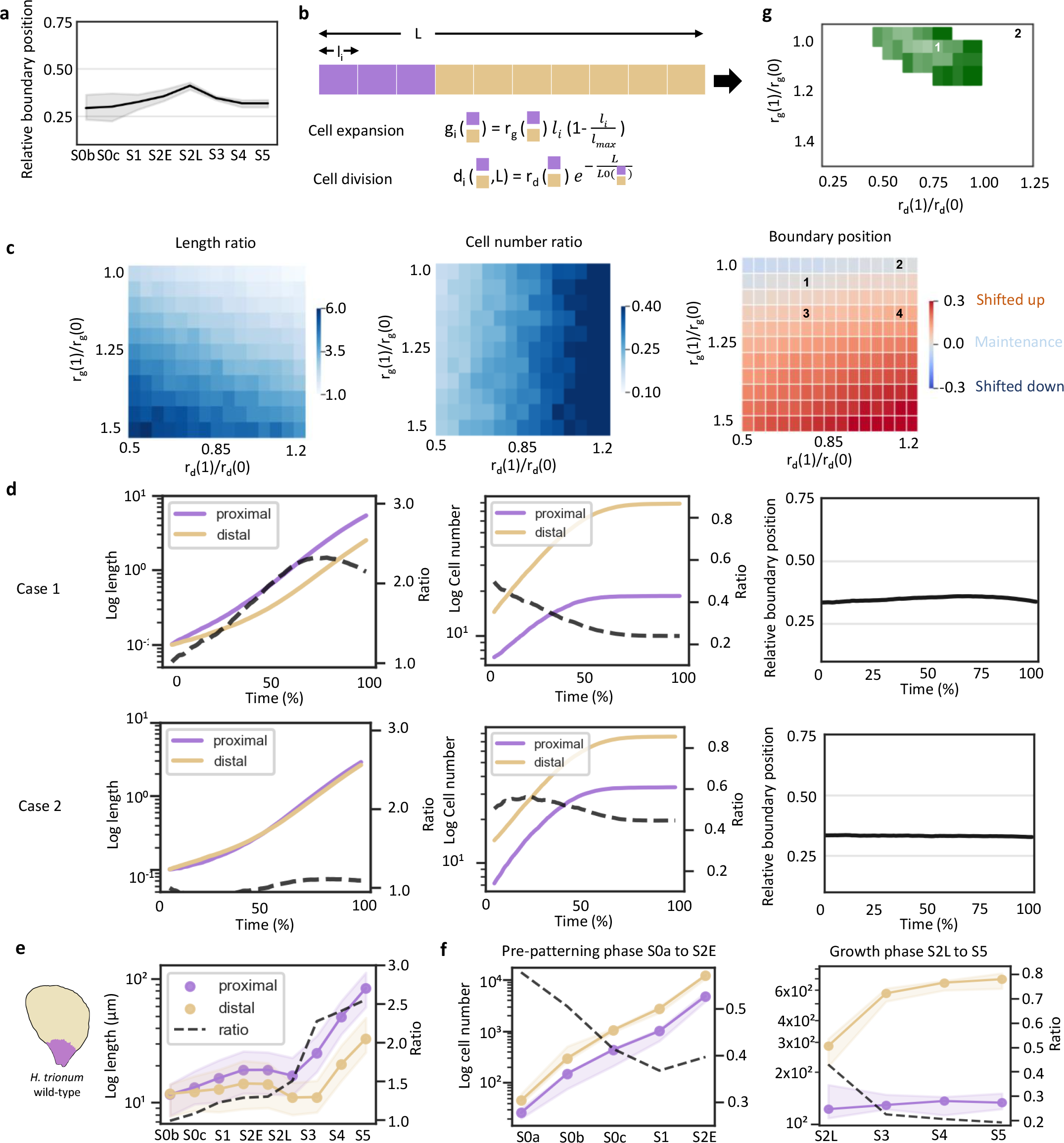
Principles governing the maintenance of the relative boundary position along the PD axis during *H. trionum* petal development. **a**, Evolution of the boundary position during *H. trionum* WT petal development. The boundary position is automatically detected as the peak of largest cells in window-averaged cell area for S0a to S2E (pre-pattern boundary) and corresponds to the pigmentation boundary, when pigmentation shifts from pink to white (bullseye boundary) for S2L to S5. **b**, Investigation of boundary maintenance principles through a 1-D cell model featuring 2 distinct cell fates (representing proximal and distal epidermis cells – See Methods). **c**, Analysis of three key observables from the simulation of the developmental process with varying ratios of growth parameters (expansion and division rates) for the two cell types: average-length ratio and number-of-cell ratio between the two cell types, and (right) deviation of boundary position from the initial 1/3^rd^ initial position (See Methods for details on calculation). **d**, Detailed plots of simulations where the model predicts boundary maintenance. **e**, Evolution of the median adaxial epidermis cell length in the proximal (below boundary) and distal (above boundary) regions during *H. trionum* WT petal development. The ratio represents the length of the proximal cells divided by that of the distal ones. **f**, Evolution of the average cell numbers in the proximal and distal regions during early stages (left, stage 0a to stage 2E) and late stages (right, stage 2L to stage 5) of *H. trionum* WT petal development **g**, An objective function representing the average percentage distance over time between simulated and experimental values for the three observables from c. The green region indicates conditions in parameter space where this distance is less than 20% (see Materials and Methods definition of the objective function). The colour shade reflects the distance, lighter shades indicating simulations closer to experimental observations and darker shades representing greater deviations.

### Principles governing boundary maintenance throughout petal growth

To mechanistically understand how bullseye proportions are conserved despite the significant growth *H. trionum* petal primordia undergo from S1 to S5 and to identify theoretical conditions that would support boundary maintenance, we used a formal model. Specifically, we aimed at investigating how the interplay between differential growth and cell division affects the maintenance of boundary position during the later phase of petal development, once the early pattern has been established. Given that our experimental focus was on boundary position along the PD axis, space was represented as a one-dimensional (1D) linear array of cells. This spatial simplification has been used in other studies where one dimension of the tissue dominates, for example when considering hormone distributions in the root ^25^. The initial state consists of a single line of 21 cells (Fig. 3b). The model incorporates the assumptions mentioned earlier: two cell types representing proximal and distal cells, with growth and division rates depending on their respective fates (Fig. 3b). Initially, the ratios of these cell types are set at the one third position from the base, and cells all have the same length. This assumes that while cell morphology is identical, proximal or distal cell fates have already been specified. Then, we simulated the late phase of petal development by reducing rates of cell division over time, while cell expansion rates also slowed down as cells reached their maximum size.

We plotted the outcomes of the model in terms of ratio of length, number of cells, and boundary position, based on the balance (proximo/distal ratio) of expansion and division rates between the two regions (e.g., ratio of 2 for distal cells divide twice as fast as proximal cells, while a ratio of 1 corresponds to an equal expansion rate between the two regions) (Fig. 3c). The results indicate that boundary position is sensitive to both cell division and expansion rates (with a higher sensitivity to growth differences) and this allowed us to single out configurations that would ensure conservation of the boundary position at one third from the petal base (Fig. 3c,d). We found that boundary maintenance occurs either when the expansion rate is higher in the proximal domain and cell division rate is higher in the distal region (Case 1, Fig. 3c,d), or when the division rate ratio is higher in the proximal domain but with similar growth rate in the two zones (Case 2, Fig. 3c,d).

To test whether one of these two scenarios could explain the boundary maintenance observed in *H. trionum* (Fig. 3a), we characterised experimentally parameters of expansion and division at later stages of development. Given the technical challenges of directly tracking petal cell growth parameters over time, we employed an approach that leverages averaged behaviours across developmental stages to reveal fundamental trends. We plotted the average cell length in both proximal and distal regions across stages (Fig. 3e), to deduce effective cell expansion rates (product of both division and expansion) and similarly, we counted cell numbers to approximate the division rates (Fig. 3f). Overall, the development of *H. trionum* petal occurs in two phases: an initial phase marked by intensive cell division activity, followed by a subsequent phase characterised by pronounced cell expansion (Fig. 3e,f). Notably, the division events occur more frequently at early stages (Fig. 3f, Supplementary Fig. 2f), and the division phase lasts longer in the distal region than in the proximal domain. The division phase continues until S3 in the distal part of the bullseye but stops earlier, around S2E-S2L, in the proximal domain (Fig. 3f and Supplementary Fig. 2f). As a result, from S3 onwards, the number of distal cells along the PD axis is five times higher than the number of proximal cells (Fig. 3f). Those proximal cells exhibit higher effective growth-rates (Fig. 3e and Supplementary Fig. 2f). Indeed, we found that from S3 onwards proximal cells are approximately twice as long as the distal cells despite cell dimensions being relatively even within the 2 domains at the start of petal development (Fig. 3e and Fig. 2b). Despite such growth disparities, the boundary position remains mostly constant throughout petal development, except for the distinct ‘bump’ at S2L (Fig. 3a). This bump can be attributed to the asynchrony in the exit from the division phase between the two regions (Fig. 3f).

In summary, we found that in *H. trionum* WT, the expansion rate is higher in the proximal domain and division events occur more frequently in the distal region. These experimental conditions (area ratios and number-of-cell ratios) do not match with the second scenario (Case 2, Fig. 3g) where a higher division rate ratio combined with similar growth rates in the two zones maintains the boundary position but leads to cell sizes and number-of-cell ratios that differ from those experimentally recorded in *H. trionum* WT (Case 2, Fig. 3d-g) however they align with the outcomes theoretically predicted by the first scenario of our simulations (Case 1, Fig. 3d-g).

Altogether, we identified that the boundary is set very early during the first stages of petal development and that local differences in cell expansion and division between the two early domains enable its maintenance, ultimately yielding the mature bullseye pattern.

### Developmental processes responsible for changes in bullseye proportions

In nature, bullseye size varies between *H. trionum* and the other members of the Trionum complex ^22^, yet the mechanisms accounting for this variation have not been investigated.

Theoretically at least two distinct mechanisms acting at different timepoints of petal development could account for changes in bullseye proportions: the position where the early pattern boundary is specified could be shifted (higher or lower) along the petal PD axis during the pre-patterning S0 phase and maintained at that position during later growth (mechanism #1), or the early boundary could remain specified at one third (pre-patterning phase unchanged) but cell expansion and proliferation could vary either side of this boundary yielding a shifted bullseye boundary in mature flowers (mechanism #2).

To test whether plants use these processes to regulate bullseye size, we took advantage of the natural diversity within the Hibiscus family and characterised pattern formation in *Hibiscus richardsonii*, a close relative of *H. trionum* that produces flowers with notably smaller bullseyes (Fig. 4a, b). In *H. richardsonii*, the pigmented area represents only 2.1% of the total petal surface, a striking contrast to the 14.5% observed in *H. trionum* (Fig. 4c). In addition to the shift in pigmentation, the bullseye is also smaller in terms of cell shape distribution. When flowers open (S5), the cell shape boundary (transition from flat striated tabular to conical smooth cells) is closer to the petal base, with the maximum cell size lying at the 0.15 position along the PD axis of the petal (Supplementary Fig. 3a). Beyond this peak, cell area declines to reach a plateau around the 0.3 to 0.4 positions before decreasing again sharply. To investigate the mechanism responsible for reduced bullseye size in this natural variant, we tracked early pattern boundary formation. From S0a to S0b, we detected no differences in cell area distribution between *H. richardsonii* and *H. trionum* petal primordia (Supplementary Fig. 3b). However, at S0c, while larger cells emerge around the one third position from the base in *H. trionum* (Fig. 2a,b), larger cells are found closer to the petal base in *H. richardsonii* (near the 0.1 position from the base) (Fig. 4d, e). By S2E, instead of occupying the one third position, the largest cells are found nested closer to the base (averaged cell area maxima observed at 0.15 position from the base). While cell areas decrease sharply after this maximum at S2E, on average the cells remain smaller than their equivalent at the one third position in *H. trionum* (around 140 μm2 vs 165 μm2 for *H. trionum*) (Supplementary Fig. 3b). Cell size then drops significantly in the distal half of the petal, following a trend already observed in *H. trionum* primordia. Thus, the reduction in bullseye dimensions that occurred on the lineage leading to *H. richardsonii* is associated with a change in cell behaviour along the petal PD axis, with the early pattern boundary specified closer to the petal base. Taken together our results suggest that the size reduction of the structural bullseye (cell shape and texture) in *H. richardsonii* follows the principle of the first theoretical mechanism outlined above, with an early boundary specified closer to the petal base during the pre-patterning phase at S0 followed by boundary maintenance during the later growth phase, as observed in *H. trionum*.

**Fig. 4.**
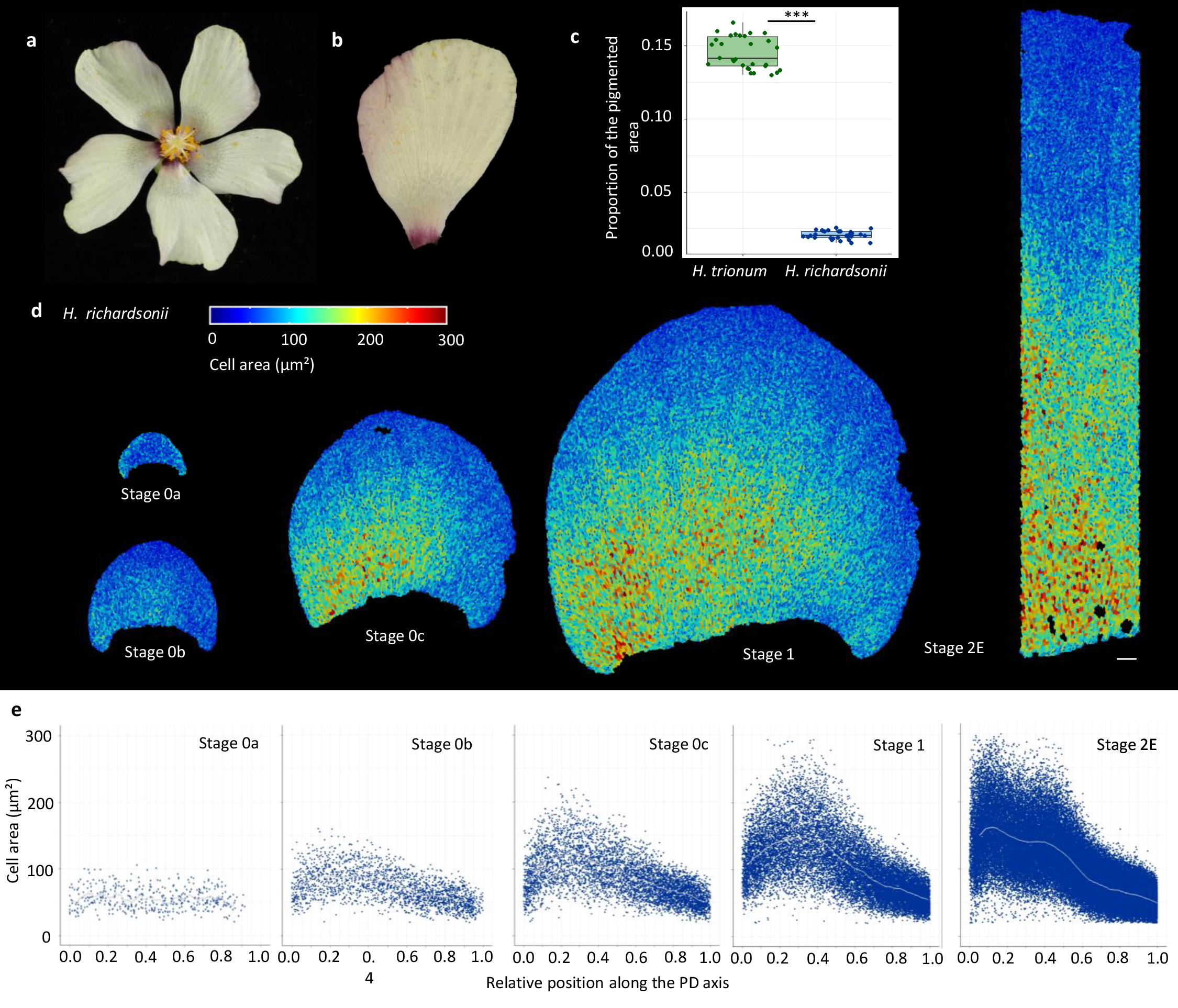
The pre-pattern boundary is specified closer to the petal base during early stages of *H. richardsonii* petal development. **a**, Flowers of *H. richardsonii* display a smaller bullseye than its sister-species *H. trionum*. **b**, Close-up view of *H. richardsonii* petals. **c**, Comparison of bullseye proportions (pigmented area/total area) in *H. trionum* and *H. richardsonii* open flowers (stage 5). n= 10 flowers and 3 petals per genotype. Statistical differences were calculated using a Shapiro-Wilk test to evaluate the normality, followed by T-test, ^***^p<0.01. **d**, Colour map of cell area across the adaxial petal epidermis of *H. richardsonii* during early developmental stages (from S0a to S2E). Scale bar, 100 μm. **e**, Cell area distribution across the PD axis of *H. richardsonii* petals. The graph consider only the central stripe of cells (20% of the petal width) for readability. Cell positions along the PD axis are relative, with 0 corresponding to the petal base, and 1 to the petal tip. Grey lines correspond to the average cell area of all replicates. n = 5 petals for each stage.

We then analysed cell behaviour across the adaxial petal epidermis of transgenic *H. trionum* lines producing flowers with a larger bullseye compared to WT (Fig. 5a, b and Supplementary Fig. 4a, b). These plants constitutively overexpress *HtTCP4*.*1 or HtTCP4*.*2* (OE *HtTCP4*.*1* and OE *HtTCP4*.*2*), transcription factors (TFs) from the TEOSINTE BRANCHED 1, CYCLOIDEA, PCF1 (TCP) family. This group of plant-specific transcriptional regulators ^26–29^ plays a pivotal role in various developmental processes, primarily by controlling cell growth, proliferation and differentiation (Nicolas & Cubas, 2016). In WT *H. trionum, HtTCP4*.*1* and its paralogue *HtTCP4*.*2* are both preferentially expressed in the distal petal region throughout petal development (Supplementary Fig. 4c). Under control of the strong constitutive 35S promoter, the ectopic activity of *HtTCP4*.*1* produced flowers with increased bullseye proportions as the pigmented area represents 25% of the total petal area instead of 14.5% recorded in WT (Fig. 5a, b and Supplementary Fig. 4a). A similar phenotype was observed when *HtTCP4*.*2* was overexpressed instead (see Supplementary Fig. 4a, b).

**Fig. 5.**
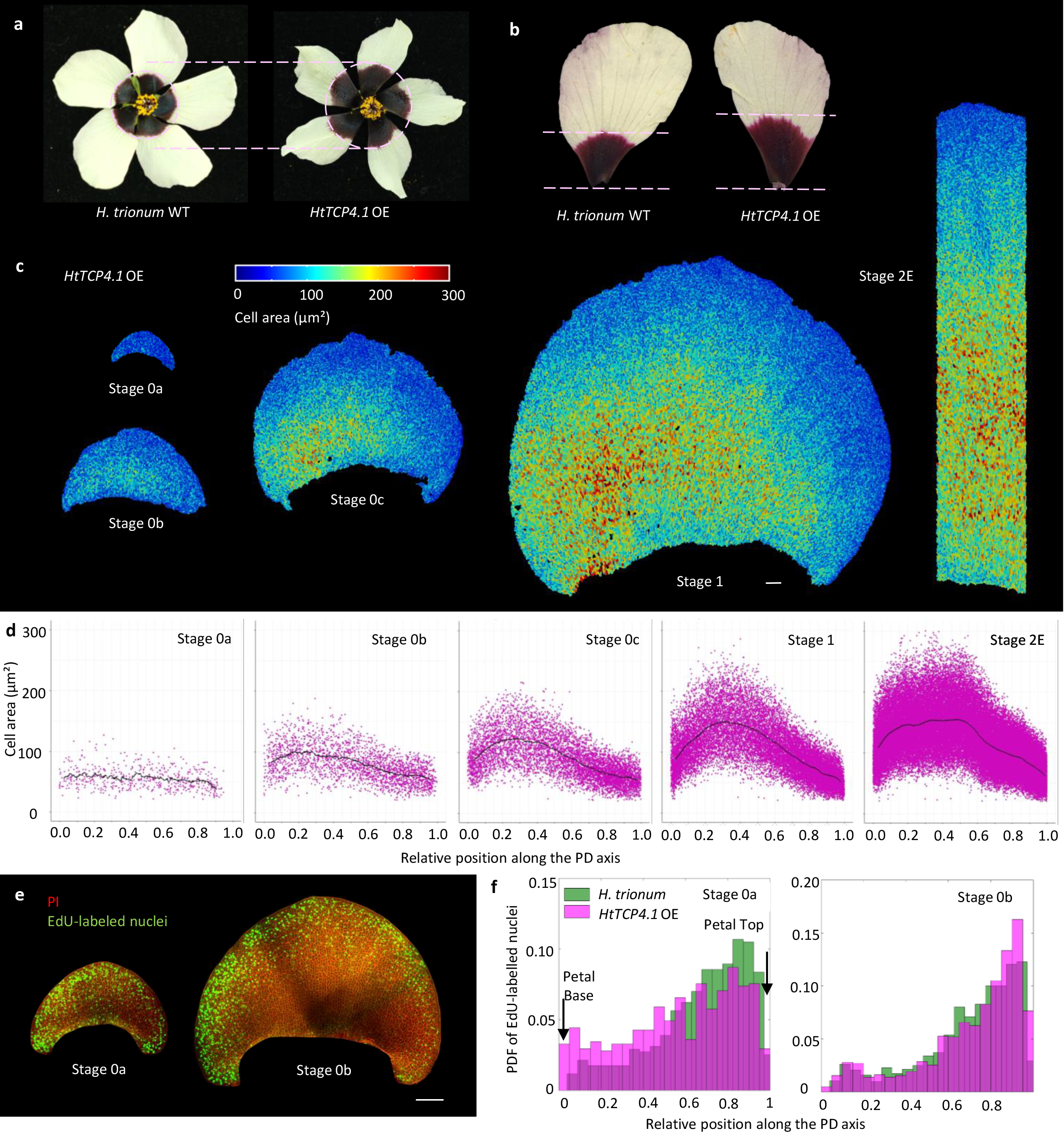
Overexpression of *HtTCP4*.*1* produces *H. trionum* flowers with a larger bullseye due to a change in spatial distribution of cell division events across the adaxial petal epidermis. **a**, Flowers of *HtTCP4*.*1* OE transgenic lines display a larger bullseye compared to WT. **b**, Close-up view of WT (left) and *HtTCP4*.*1* OE (right) petals. The pink dotted lines indicate the distance between the petal basis and the bullseye boundary. **c**, Colour map of cell area across the adaxial epidermis of *HtTCP4*.*1* OE petals during early developmental stages (from S0a to S2E). Scale bar, 100 μm. **d**, Cell area distribution across the PD axis of *HtTCP4*.*1* OE petals. The graphs consider only the central stripe of cells (20% of the petal width) for readability. Cell positions along the PD axis are relative, with 0 corresponding to the petal base, and 1 to the petal tip. Black lines correspond to the average cell area of all replicates. n = 5 petals for each stage. **e**, Distribution of cell division events across the adaxial epidermis of S0a and S0b *HtTCP4*.*1* OE petals. Newly synthesized DNA is labelled using fluorescently labeled nucleotide analog 5-ethynyl-2-deoxyuridine (EdU) (green) and plasma membranes are stained with propidium iodide (red). Scale bar, 100 μm. **f**, Probability density function (PDF) of the EdU-labeled nuclei along the PD axis of *HtTCP4*.*1* OE S0a petals compared to *H. trionum* WT (stripes corresponding to 20% of the petal width and centred along the PD axis were analysed, see Supplement Fig. 4e). n = 5 petals for each stage.

We found no significant difference in cell area distribution along the PD axis during the pre-patterning phase (from S0a to S1) in OE *HtTCP4*.*1* petals compared to WT (Fig. 5c, d): cell area is uniform across the epidermis of both genotypes at S0a and an early croissant-shaped pattern emerges at S0b resulting in a right-left asymmetry (Fig. 5c). At S0c, cell area peaks at one third of the petal length from the base, as in WT (Fig. 5d). By S1, the maximum cell size at the peak position was similar for both OE *HtTCP4*.*1* and WT (Fig. 2b and 5d). This suggests that the mechanism leading to a larger bullseye in transgenic individuals constitutively overexpressing *HtTCP4*.*1* is not a change in the specification of the early boundary during the pre-patterning phase. At S2E, we observed a higher proportion of large cells in the proximal region of the petal in OE *HtTCP4*.*1* petals compared to WT (Fig. 5c). Those petals displayed a plateau of larger cells starting closer to the petal base and expanding beyond the one third landmark of the petal length (Fig. 5d), a trend that can also be observed later along the PD axis of the mature petal at S5 (Supplementary Fig. 5c). While both WT and transgenic petals reached the same maximum average cell size at S2E (Fig. 2b and 5d), the proportion of cells reaching this size is significantly increased when *HtTCP4*.*1* is constitutively overexpressed. No significant differences were observed in the distribution of cell division events from S0b to S1 between OE *HtTCP4*.*1* and WT petal primordia (Fig. 5e, f and Supplementary Fig. 4f, g). However, at S0a, instead of the active proliferation zone being restricted to the distal part of the petal, as observed in WT, EdU-labelled nuclei were detected across the entire adaxial epidermis of OE *HtTCP4*.*1*, including the proximal region (Fig. 5e, f and Supplementary Fig. 4e), indicating cell proliferation is more uniform across the adaxial petal epidermis when *HtTCP4*.*1* is constitutively overexpressed.

Altogether, these findings indicate that the larger bullseye in *HtTCP4*.*1* OE is not due to a shift in the specification of the early pattern boundary along the petal PD axis during the S0 phase but might instead be due to an increase in cell proliferation at the petal base. These additional basal cells, which have the fate of the proximal domain, are programmed to grow more than the distal ones, and this could explain the higher proportion of larger cells at the petal base, ultimately resulting in a larger bullseye.

To test this hypothesis further, we first used our formal model to identify possible conditions that could interfere with maintenance of the early boundary (Fig. 3c). We found several theoretical configurations that could account for an upward shift of the final bullseye boundary in mature flowers: first, increasing the growth rate ratio shifts the boundary position upward, while area ratios and number of cell ratios remain similar to those observed in WT petals (case 3, Fig. 3c and 6 a). Increasing both the division rate and growth rate ratio (greater growth in the proximal region) shifts the boundary position upward and alters the number-of-cell ratios and area ratios (case 4, Fig. 3c, Fig. 6a).

**Fig. 6.**
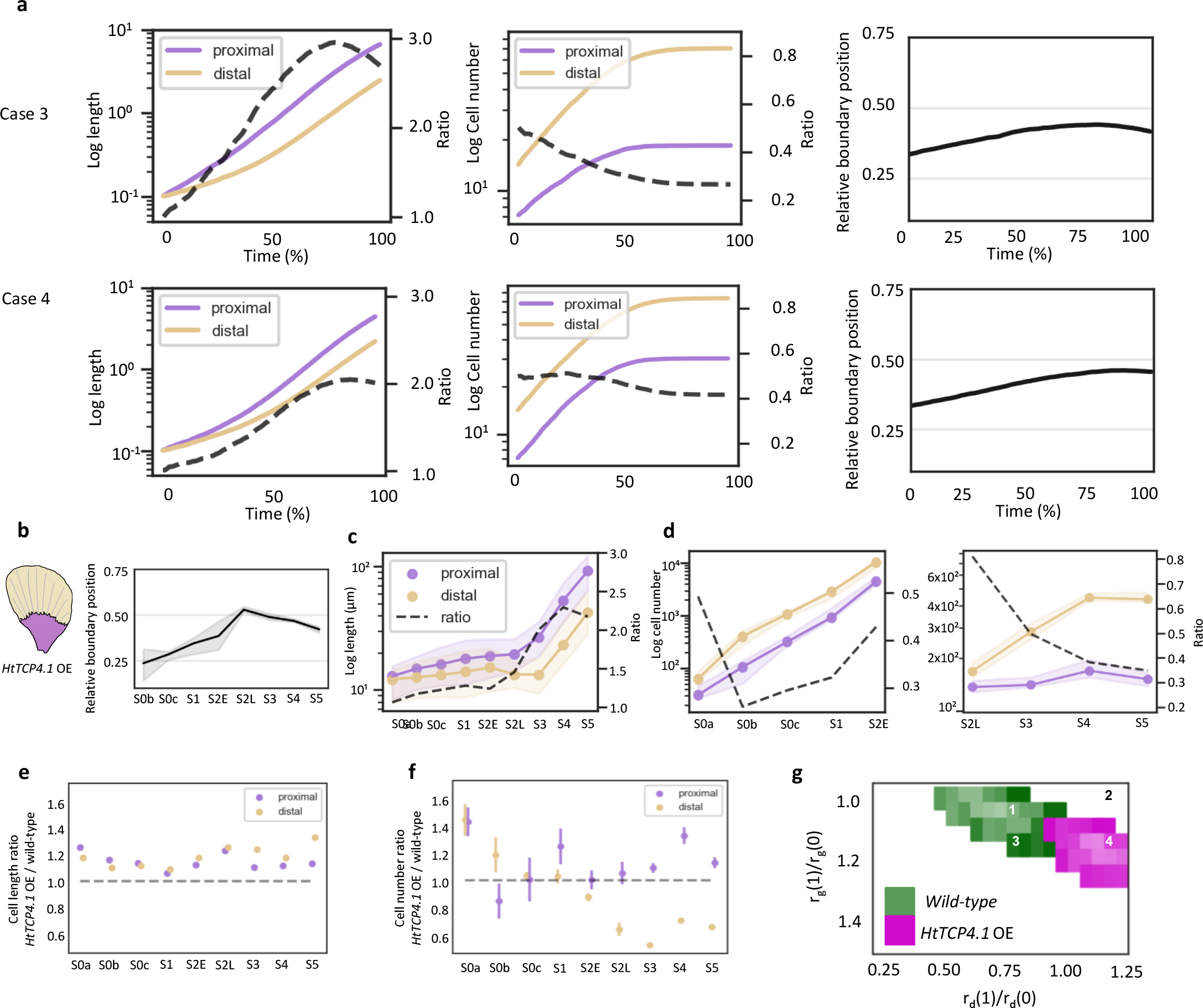
Evolution of the boundary position and epidermis cell features during petal development in transgenic lines overexpressing *HtTCP4*.*1*. **a**, Detailed plots of simulations where the model predicts a shift in the boundary position. **b**, Evolution of the boundary position during *HtTCP4*.*1* OE petal development. The boundary position is automatically detected as the peak of largest cells in window-averaged cell area for S0a to S2E (pre-pattern boundary) and corresponds to the pigmentation boundary, when pigmentation shifts from pink to white (bullseye boundary) for S2L to S5. **c**, Evolution of the median adaxial epidermis cell length in the proximal (below boundary) and distal (above boundary) regions during *HtTCP4*.*1* OE petal development. The ratio represents the length of the proximal cells divided by that of the distal ones. **d**, Evolution of the average cell numbers in the proximal and distal regions during early stages (left, stage 0a to stage 2E) and late stages (right, stage 2L to stage 5) of *HtTCP4*.*1* OE petal development. **e**, Comparison of median cell length in *HtTCP4*.*1* OE vs. WT in the proximal and distal regions during *H. trionum* petal development. **f**, Comparison of average cell numbers in *HtTCP4*.*1* OE vs. wild-type in the proximal and distal regions during *H. trionum* petal development. **g**, An objective function representing the average percentage distance over time between simulated and experimental values for the three observables. Coloured regions (green for WT and purple for *HtTCP4*.*1* OE) indicates conditions in parameter space where this distance is less than 20% (see Materials and Methods definition of the objective function). The colour shade reflects the distance, lighter shades indicating simulations closer to experimental observations and darker shades representing greater deviations.

Next, to test whether petal morphogenesis in the *HtTCP4*.*1* OE transgenic line follows one of the configurations predicted by our theoretical model, we extended our examination of cell behaviour to the later phase of petal morphogenesis (S2L to S5). Although the early pattern boundary in the *HtTCP4*.*1* OE transgenic line is established at one third of the petal length (Fig. 5c,d) as in WT, our analysis of subsequent developmental stages reveals that the relative position of the bullseye boundary then rises to 0.5 at S2L before stabilising around the 0.4 position at later stages (Fig. 6b). We found that petals of *HtTCP4*.*1* OE follow a morphogenetic process similar to the WT: a phase of intense cell division followed by a period of cell expansion, with the distal cells exhibiting higher division rates during early development and proximal cells having higher effective growth rates (Fig. 6c). However, while the evolution of the cell area ratio (proximo/distal) is comparable to the one of WT *H. trionum*, the cell number ratio (proximal/distal) reaches a plateau at around 35% at S4, rather than 20% at S3 as observed in WT (Fig. 6d). While petals of both genotypes have similar length, a detailed comparison reveals that both proximal and distal cells are overall longer in the *HtTCP4*.*1* OE transgenic line (Fig. 6e). Additionally, there are more proximal cells but fewer distal cells (Fig. 6f) when *HtTCP4*.*1* is constitutively overexpressed, resulting in an overall lower cell number along the petal PD axis, accounting for the overall maintenance of petal length.

These experimental conditions (area ratios and number-of-cell ratios) do not match with the third scenario (Case 3, Fig. 6a,g) because it shifts the boundary position but leads to cell sizes and number-of-cell ratios that differ from those experimentally recorded in *HtTCP4*.*1* OE transgenic line, however they align with the outcomes theoretically predicted by the fourth scenario of our simulations (Case 4, Fig. 6a,g).

Taken together, our results suggest that the increased bullseye in OE *HtTCP4*.*1* follows the second theoretical mechanism outlined earlier, where the early boundary is specified one third from the petal base, as in WT but with subsequent changes in growth (increase in division and expansion), shifting the boundary position upwards and producing flowers with a larger bullseye.

### Bumblebees can discriminate targets solely based on bullseye size

Petal patterns are believed to enhance flower attractiveness and help visiting animals form a search-image to identify targets effectively ^2^. To investigate whether pollinators can discriminate between Hibiscus bullseyes of different sizes and exhibit any innate preference for specific pattern proportions, we conducted experiments with naïve buff-tailed bumblebees (*Bombus terrestris*), known to pollinate Australian and New Zealand plants ^30,31^. We created 3D-printed discs that mimicked the colour patterns of Hibiscus bullseyes. These discs were designed to match the dimension of open flowers and differed only in the size of their pigmented area, replicating the bullseye proportions of *H. trionum* WT (medium bullseye), *HtTCP4*.*1* OE (large bullseye) or *H. richardsonii* (small bullseye) flowers with pigmented area accounting for 16%, 36% and 4% of the disc surface respectively (Fig. 7a). This allowed us to assess bumblebees’ response to variations in bullseye size when no other stimuli are present.

**Fig. 7.**
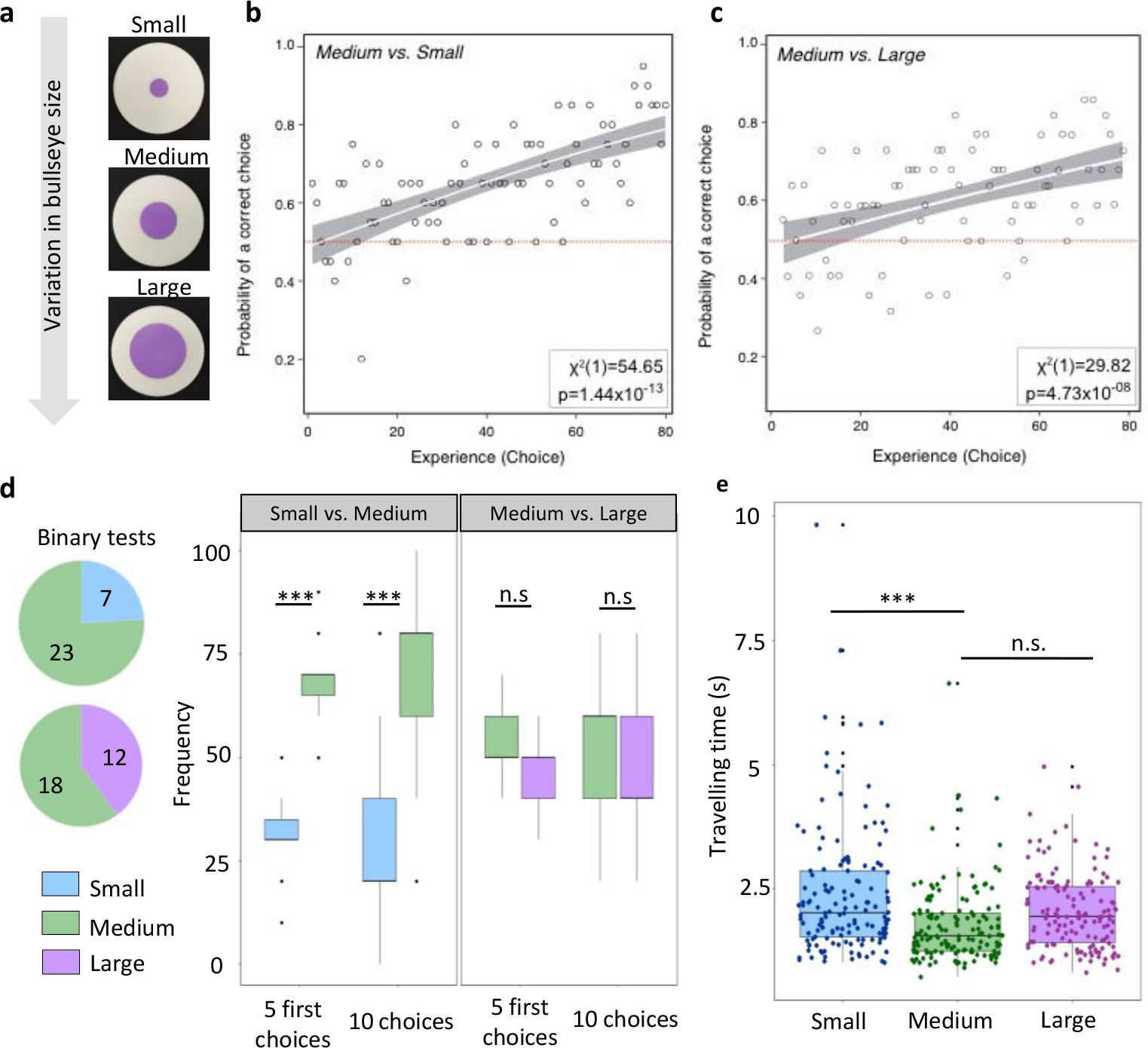
Bumblebees (*Bombus terrestris*) responses to varying bullseye sizes. **a**, Epoxy discs featuring small (*H. richardsonii*-like), medium (*H. trionum* wild type-like) and large (*HtTCP4*.*1* OE-like) bullseyes. Purple centres represent 4%, 16% and 36% of the total area, respectively. **b**, Learning curve of 20 individuals choosing between discs with small or medium bullseye sizes. Empty circles depict the mean proportion of bees choosing correctly for 80 successive choices. The white curve represents the fitted binomial logistic model, with grey shading indicating 95% confidence intervals on the fitted response. The χ2 statistic (the number in brackets indicates d.f.) and P value for the likelihood ratio test (assessing whether foragers can learn) are provided. **c**, Learning curve of 22 individuals choosing between discs with medium or large bullseye sizes. Similar annotations as in (b) are included. **d**, Preference tests experiments. See statistics in Supplementary Fig. 6. Binary test – number of naïve bumblebees choosing to first land on a disc with a small vs. medium bullseye (top pie chart) or on a disc with a medium vs. large bullseye. Bumblebees showed a statistically significant preference for the medium bullseye size compared to the small one. n=30 bumblebees for binary tests. 10 choice tests – (left) when the first five choices or first 10 choices were considered, bumblebees showed a statistically significant preference for the medium bullseye size compared to the small one. (right) when the first five choices or first 10 choices were considered, bumblebees showed no significant preference for the medium *vs*. the large bullseye (one sample t-test). n=15 bumblebees. **e**, Distribution of individual travel time between discs for the three bullseyes size. n=15 bumblebees for each bullseye size, each bumblebee flew 10 times between each disc type. Each dot corresponds to the flying time between two discs for each bee on each travel path.

To familiarise bumblebees with our foraging setup, we initially trained them to feed on black artificial discs containing a 15% sucrose solution. We conducted differential conditioning experiments to investigate their ability to distinguish between different bullseye sizes by assessing whether they could learn to associate specific proportions to the presence or absence of a reward. First, we randomly arranged five small bullseye discs (resembling *H. richardsonii*-like) and five medium-sized ones (resembling *H. trionum*-like)) in the flight arena. We assessed the behaviour of 20 bumblebees, with 10 offered a 20% sucrose solution (reward) on the medium-sized bullseye flower, and water (neutral) on the small-sized one, and the other 10 offered the opposite combination. Discs were refilled, and their positions randomised throughout the experiment. After 80 visits, bumblebees chose the rewarding discs (sucrose solution, correct choice) much more frequently than at the beginning of the experiment [learning curve (χ^2^ = 54.65, *P* < 0.001) (Fig. 7b)]. Initially, individuals visited the first ten discs randomly (55% of correct choices, 95% confidence interval [47.4%-63.6%]) but after 80 visits, the probability of making a correct choice (reward) significantly increased (82% of correct choices, 95% confidence interval [78.0%-87.0%], P = 5.57e-12). This indicates bumblebees can discriminate between small and medium bullseyes solely based on size. When examining individual behaviours, we noticed that five out of 20 individuals already displayed a preference for the medium-sized bullseye (*H. trionum*-like) at the beginning of the experiment (70-90% correct choices during the first ten visits), and the reward was consistently associated with medium bullseye in these cases (Supplementary Fig. 6a). Only two individuals showed no evidence of learning to distinguish between the two bullseyes (probability of correct choice during the last 10 visits = 0.6), and in both cases, the reward solutions were associated with the smaller, *H. richardsonii*-like pattern. This suggests that naïve bumblebees may have an innate preference for the *H. trionum* WT bullseye dimensions over the smaller, less conspicuous *H. richardsonii*-like pattern.

To determine whether bumblebees could differentiate between medium and larger bullseye, we conducted a similar experiment using 3D-printed discs featuring medium (*H. trionum*-like) and large (*HtTCP4*.*1* OE-like) pigmented bullseyes (Fig. 7a). Analysing the collective behaviour of all 22 individuals tested (Fig. 7c), our results indicate that bumblebees were capable of learning which type of bullseye was associated with a reward (χ2= 29.82, P < 0.001). Bumblebees randomly chose which disc to visit during their first ten visits (50.5% of correct choices, 95% confidence interval [44.4%-56.5%]) but after 80 visits, individuals chose correctly (rewarding disc) almost three out of four times (72.7% correct choices, 95% confidence interval [65.9%-79.6%], P = 8.39e-07). However, learning to discriminate between medium and large bullseye sizes appeared to be more difficult for bumblebees than distinguishing between medium and small patterns. When analysing individual behaviour, half of the 22 bumblebees were able to associate bullseye size with presence/absence of a reward (χ2= 54.79, P < 0.001; Supplementary Fig. 6b), while the other half did not (χ2= 1.74, P >0.5; Supplementary Fig. 6c). Altogether, these results suggest that, on average, it may be more challenging for bumblebees to discern the size difference between a medium (*H. trionum* WT pattern) and a large bullseye (*HtTCP4*.*1* OE pattern) compared to the medium *vs*. small bullseye combination. However, for those individuals that successfully differentiate medium patterns from large ones, their performances matched those of bumblebees asked to distinguish between medium and small bullseyes (compare Fig. 7b with Supplementary Fig. 6b).

### Bumblebees prefer the *H. trionum* WT bullseye proportions over those of its close relative *H. richardsonii*

Next, we used a binary choice experiment to test whether naïve bumblebees display an innate preference for specific bullseye proportions. Two equally rewarding discs (20% sucrose solution), one displaying a small bullseye and the other presenting a medium pattern, were positioned equidistant from the hive entrance, and a single naïve forager was allowed to enter the arena, with its first choice recorded. Out of 30 bumblebees, 23 chose to land first on the medium-size bullseye disc (Fig. 7d, Supplementary Fig. 6d). To evaluate whether this preference persisted during a foraging bout, we randomly placed five artificial discs of both bullseye types across the arena, all containing the rewarding solution. We recorded the first 10 choices made by 15 naïve bumblebees, refiling and changing the flower position as the individuals foraged. Whether considering the first five or first ten choices, bumblebees consistently preferred the medium-sized bullseye (*H. trionum* WT) over the smaller one (*H. richardsonii*). Specifically, the trionum-like discs were chosen 7 out of 10 times (Fig. 7d, Supplementary Fig. 6d). We repeated this experiment using medium (*H. trionum* WT) and larger (*HtTCP4*.*1* OE) bullseyes. In this case, regardless of whether we considered the binary choice, first five choices, or all 10 choices, we could not detect any statistically significant preference for either of the two bullseye sizes (Fig. 7d, Supplementary Fig. 6d).

### Enlarged bullseye size enhances flower detection

To investigate the possible impact of bullseye size on flower detectability we recorded the time individuals took to move from one target to the next (foraging speed) for each of the three types of pattern dimensions. We found that bumblebees flew significantly faster (P= 0.0023) between artificial flowers displaying a medium-sized bullseye (*H. trionum* WT) compared to those foraging on discs with a smaller pattern (*H. richardsonii*) (Fig. 7e). However, no significant difference in mean travel time was observed when comparing large bullseyes (*HtTCP4*.*1* OE) to medium-sized ones (P=0.23), consistent with our previous findings indicating bumblebees may find distinguishing between those two bullseye sizes challenging.

Overall, our data indicate that foragers can discern between targets solely based on bullseye size differences and use pattern dimensions as a reliable cue to identify rewarding flowers. Our findings also demonstrate that bullseye size directly impacts flower detectability and that buff-tailed bumblebees exhibit a strong innate preference for *H. trionum* bullseye over the smaller pattern of its close relative, *H. richardsonii*.

## Discussion

Our analysis of developing *H. trionum* petal primordia revealed that cell behaviour across the adaxial epidermis is pre-patterned, characterised by regional differences in cell expansion and division along the base-to-tip axis (PD axis) long before a visible bullseye emerges. Following an initial stage of uniform behaviour (S0a), the petal distal domain mainly grows through cell division, while the size of the proximal region increases predominantly through cell expansion. From Stage 0b onwards, the largest cells are concentrated in a region invariably positioned one third from the petal base. These cells could represent the first cells to initiate differentiation across the petal adaxial epidermis, becoming anisotropic by elongating preferentially along the PD axis. Notably, this landmark also corresponds in later stages to the transition point between pigmented and non-pigmented cells that characterise the final bullseye boundary. Hence, the pre-patterning phase may already specify bullseye boundary cells early on, influencing pattern proportions. However, further investigations are needed to determine whether the largest cells emerging during the pre-patterning phase are indeed the first to differentiate and act as progenitors for the bullseye boundary cells. Regardless of whether the early pattern boundary yields the final bullseye boundary, our results show that the partitioning of the adaxial epidermis into subdomains during early petal development has a significant influence on the emergence of distinct cell behaviours in neighbouring regions of the epidermis tissue. Indeed, an early pattern boundary specified closer to the petal base is associated with the production of a smaller bullseye in *H. richardsonii*. Interestingly, the pigmented area in *H. richardsonii* is even smaller than the proximal domain. This indicates that although the shift in early boundary specification we uncovered along the petal PD axis is sufficient to produce the smaller structural bullseye (i.e., flat tabular striated cells cover a smaller portion of the total petal area), additional changes affecting gene(s) controlling pigment production, acting downstream of those controlling the pre-patterning process, must also have occurred along the lineage leading to *H. richardsonii* during evolution.

While our study focuses on *H. trionum* and its closest relative, pre-patterning the petal epidermis along the PD axis likely represents a general mechanism shared by a multitude of species. For instance, a Turing-like process was recently proposed to produce the spotted patterns on the ventral petals of *Mimulus lewisii* and *Mimulus guttatus*, both Monkey flowers. The suggested mechanism relies on a tug-of-war between two transcription factors from the MYB family: NEGAN (NECTAR GUIDE ANTHOCYANIN), an activator of anthocyanin pigment production ^16^, and RTO (RED TONGUE), a repressor of NEGAN ^11^. This model is particularly elegant as it adheres to the principle of self-organisation ^32^ and does not require the existence of an early pattern. In natural variants or knockout lines where RTO activity is absent, the spots are replaced by uniform pigmentation. However, this pigmentation does not extend to the entire petal epidermis but remains confined to the distal part of the petal, resembling a red tongue. This suggests that, in addition to the spotted phenotype generated by a Turing-like system, the petal epidermis is also compartmentalised into distinct domains along the PD axis. The absence of RTO activity removes pigment production repression within one of those domains (the distal region) and renders the existence of petal compartments apparent. To further explore this hypothesis, it would be valuable to investigate whether the identity of the future yellow proximal and red distal regions of the Mimulus petal are also specified very early in development during a pre-patterning phase similar to what we uncovered in Hibiscus.

Pre-patterning may represent an ancestral process plants employed to specify cell fate along the different axes of their lateral organs long before flower originated. Indeed, a pre-pattern mechanism has been proposed to play a role in establishing abaxial – adaxial polarity in leaves, with spatial information provided by the activities of REVOLUTA, AS2 and KANADI1 across the shoot apical meristem, positioning the adaxial auxin response ^33,34^. From an evolutionary viewpoint, petals can be viewed as modified leaves, thus it will be important to identify the molecular players orchestrating petal patterning and test whether those differ from the agents responsible for leaf patterning.

While the mature bullseye at S5 exhibits clear bilateral symmetry, the early development of the petal epidermis surprisingly reveals a right-left asymmetry. Initially, larger cells first emerge near the attachment point to the floral structure and the ovary base on one side of the petal. Notably, the thickness of the petal base is uneven, and the side where the early pattern initiates consistently aligns with the thicker side, likely having a stronger connection to the rest of the floral structure. This suggests that an upstream positional signal, produced externally during the early phase of bud development, could be responsible for pre-patterning the petal epidermis, with the cells closest to the source of this signal being the first to modulate their behaviour. Not only does the early pattern arise from one side, but the emergence of pigmentation is also asymmetric. Colouration initially appears as two dots on either side of the petal attachment point at S1 (Fig. 1c), yet the pigmented mark associated with the thicker side of the petal base always forms earlier and often appears larger. This reinforces the idea that pre-patterning of early cell behaviour and the development of bullseye features are closely intertwined.

Flower patterns come in diverse types (stripes, spots, bullseye, etc) but also exhibit variations in dimensions. Here, we explored the mechanisms contributing to the variation in bullseye proportions. We conducted a comparative analysis in flowers with different bullseye sizes: a transgenic line overexpressing *HtTCP4*.*1*, resulting in a larger bullseye compared to WT *H. trionum*, and a close relative of *H. trionum, H. richardsonii*, which exhibits a significantly smaller bullseye. In *H. richardsonii*, the peak of larger cells first become apparent at S0c, positioned around 0.15-0.2 from the petal base (in contrast to 0.3 for *H. trionum*). This indicates a downward shift of the pre-pattern boundary along the petal PD axis. From S2 onwards, the peak of larger cells remains closer to the base (around the 0.1-0.2 position), aligning with the position of the boundary typical of the smaller bullseye of *H. richardsonii*. These findings support the idea that (i) the early pattern could determine the final bullseye dimensions, and (ii) early pattern proportions can be maintained while the petal grows to S5, allowing the bullseye to scale up along with flower size. Contrastingly, early boundary emergence from S0a to S1 in the *HtTCP4*.*1* OE line is similar to WT implying the significant increase in bullseye dimensions observed in mature flowers is not due to an early shift in early pattern boundary positioning. Instead, both computational simulations and experimental observations support the idea that the larger bullseye is due to later changes in growth, acting as pattern-modifiers. At S2, cellular behaviour diverges from WT, with a higher proportion of large cells spreading around the 0.2 to 0.5 position. Unlike WT petals where cell division and DNA replication are mainly restricted to the distal region, cell proliferation events occur across the entire S0a petal adaxial epidermis when *HtTCP4*.*1* is overexpressed. This suggests that the larger bullseye observed in the *HtTCP4*.*1* OE line results from an excess of cell proliferation at the petal base early in development (S0a), positioning more cells to acquire the fate of the proximal domain (i.e., change in initial conditions). Proximal cells, programmed to grow more than the distal ones during the differentiation process, lead to larger cells at the petal base (i.e., change in growth), ultimately resulting in a larger bullseye. Taken together, these results illustrate how local variations of growth and cell proliferation on either side of the early pattern boundary can act as a robust mechanism for modulating pattern proportions and thereby regulate the dimensions of the final bullseye. Thus, partitioning the petal epidermis into subdomains not only plays a role in controlling cell fate specification spatially but it also constitutes an effective system for autonomously regulating growth in two neighbouring domains of the epidermis tissue, where DNA replication, cell division and expansion are independently controlled.

Hormonal crosstalks, especially the balance between auxin and cytokinin are central to the patterning of Arabidopsis roots, ovary and grasses leaves ^35–37^. Considering the well-known roles of both hormones in regulating cell proliferation and expansion, it is likely that plant hormones also contribute to the control of bullseye dimensions. For instance, TCP4 in Arabidopsis has recently been shown to promote auxin synthesis during development via transcriptional activation of *YUCCA5* expression ^38^. *HtTCP4*.*1* is preferentially expressed in the distal portion of the *H. trionum* petal, where most cell division events occur, and ectopic overexpression of *HtTCP4*.*1* is sufficient to induce excessive cell proliferation in the proximal region at S0a. These observations suggest that HtTCP4.1 participates in setting the bullseye dimension by promoting cell proliferation in the distal domain and that manipulating its spatiotemporal expression constitutes a means to modify bullseye proportions using a pattern modifier process. Whether HtTCP4.1 activity in Hibiscus relies on local activation of auxin production will need to be tested in further studies.

We found that bumblebees can effectively distinguish between small and medium bullseyes mimicking those of *H. trionum* and *H. richardsonii*, respectively, based on size differences only. Even without a strong incentive (without using quinine hemisulfate salt solution as punishment ^19^, but rather a neutral solution of water against rewarding sugar solution), bumblebees successfully discriminated between the two bullseye sizes, indicating that they can easily detect the difference. Preference and binary choice tests further revealed that buff-tailed bumblebees have an innate preference for *H. trionum*-like bullseye size over the smaller pattern of its relative *H. richardsonii*. Foraging tests also showed that bumblebees could detect targets faster when a medium rather than a small bullseye was present on the discs. Further investigations are required to determine whether this preference holds in a more realistic context, when additional elements like UV or scent might compensate for the reduction in pigmentation, potentially affecting overall attractiveness. However, our results indicate that flowers with reduced bullseye could be discriminated against when growing alongside flowers with larger patterns. Notably, *H. richardsonii* is classified as a “Vulnerable” species in Australia and as “Threatened/Nationally Critical” in New Zealand ^22^, with populations declining in their natural habitat. While the exact causes remain uncertain and are likely to be multiple, the reduction in bullseye size may contribute to a decline in pollinator attraction. Importantly, our behavioural experiments have focused on buff-tailed bumblebees, and the response of other foraging insects might differ. One intriguing hypothesis is that a change in bullseye dimensions could mediate a change in pollinator type. Whether a change in pattern proportions can lead to reproductive isolation and promote speciation are open questions that will certainly necessitate field investigations.

To conclude, our study highlighted that the establishment of a pre-pattern is a key feature of Hibiscus petal development. Events affecting the patterning process itself early in development (modification of the early pattern boundary position) or processes acting as pattern modifiers at later stages (local change in growth/cell division either side of the boundary) represent two distinct mechanisms equally able to produce variations in bullseye proportions (Fig. 8). Such modifications in pattern dimensions hold crucial biological importance, as buff-tailed bumblebees can distinguish flowers based on bullseye size only and exhibit an innate preference for medium-sized patterns over smaller ones. What genetic bases account for differences in bullseye size between the two sister species and whether such a change in pattern dimension contributed to reproductive isolation and speciation represent interesting venues for future research.

**Figure 8.**
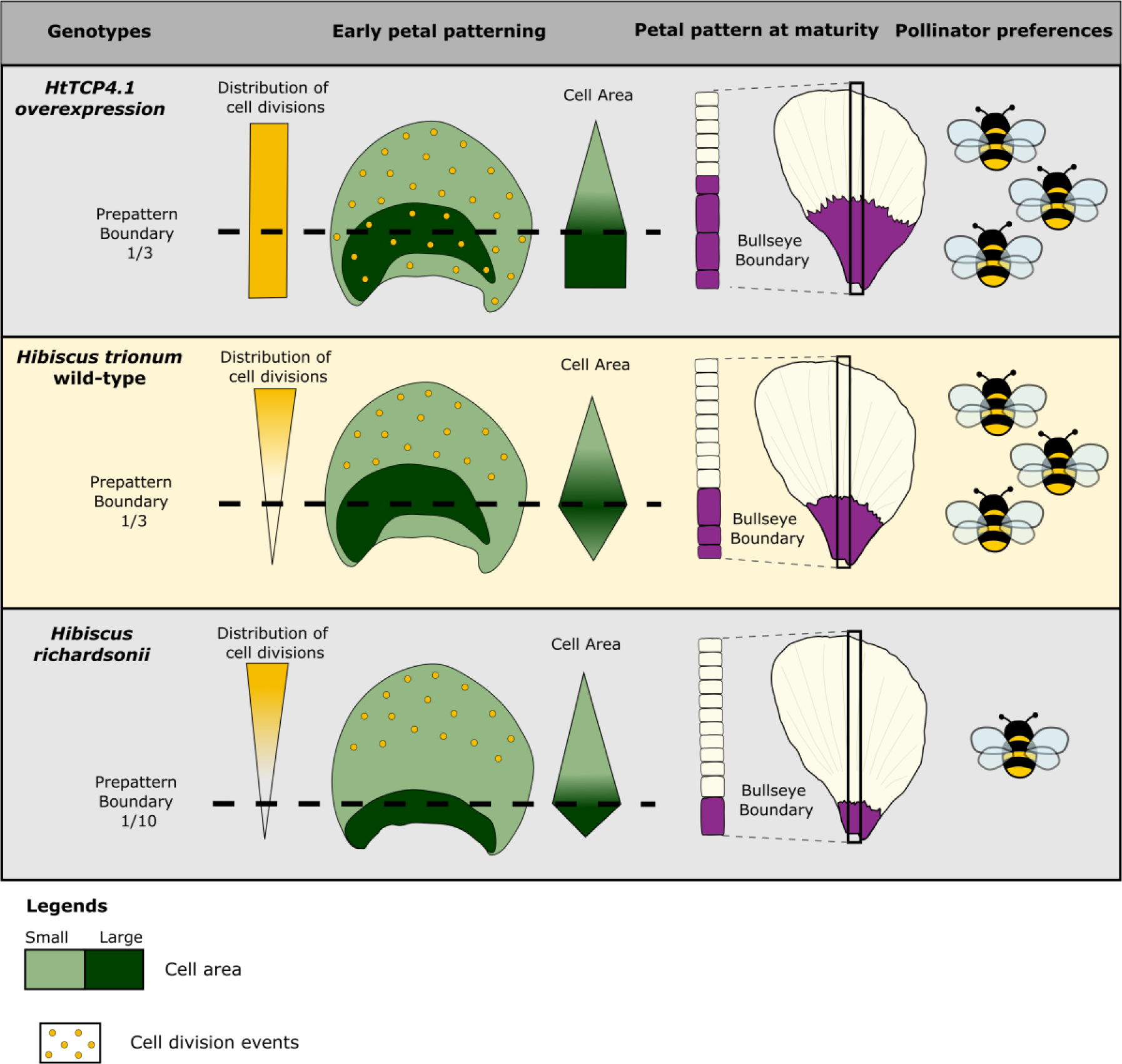
Summary of processes involved in setting up bullseye pattern proportions and its impact on bumblebee behaviour.

## Material and Methods

### Plant material

*H. trionum* L. seeds were originally obtained from Cambridge University Botanic Garden; *H. richardsonii* seeds [Mayor Island (Tuhua), New Zealand - Voucher AK251841] were kindly supplied by Prof. B.G. Murray ^39^. Wild-type *H. trionum* and *H. richardsonii* plants and *35S::HtTCP4*.*1* or *35S::HtTCP4*.*2* transgenic *H. trionum* lines were grown under glasshouse conditions on a 16h:8h, light: dark photoperiod at 23°C in Levington’s M3 (UK) compost.

### Production of the HtTCP4.1 and HtTCP4-like 2 OE lines

Gibson assembly ^40^ and primers oEM250-F to oEM257-R were used to insert the full-length coding sequence of *HtTCP4*.*1* into a modified pGREEN II vector backbone containing a double 35S promoter (pEM110), yielding the plant expression vector pEM105. The coding sequence of *HtTCP4*.*2* was introduced into the same pEM110 backbone using the BanH1 and HindIII entry sites and primers oMY049-F (ATAAGCTTTAATGGGGGACAGCCAC) and oMY047-R (AGGGATCCCTTCAATGGTGAGAATCGGACGA), yielding the plant expression vector pMY40. The coding sequences of *HtTCP4*.*1* and *HtTCP4*.*2* have been deposited in GenBank under the accession number OR908924 and OR985928, respectively. A transgenic *H. trionum* lines overexpressing constitutively *HtTCP4*.*1* or *HtTCP4*.*2* were obtained using pEM105 and pMY40 respectively and Agrobacterium-mediated transgene delivery followed by tissue culture to induce callus production and plant regeneration, following the protocol of ^18^.

### Imaging and image analysis

#### Distribution of the cell features across the petal from S0a to S2E using confocal microscopy

*H. trionum, HtTCP4*.*1* OE and *H. richardsonii* petals were dissected from S0a to S2E and mounted on a petri dish with double-sided tape. 3D depth-composition images of each petal were acquired using a Keyence VHX-7000 digital microscope at 100 to 300X magnification. Petals were stained with 0.01 μg.μL^-1^ FM1-43 (Thermo Fisher Scientific) for 15 min to 5 hours depending on their stages (respectively S0a and S2E) and incubated in the dark. Petals were washed twice with water before imaging using a 20X water-dipping objective on a LSM880 confocal microscope (Zeiss, Germany), excitation at 514 nm and emission filters set to 537-622 nm. For S0a to S1 primordia, the whole petal was imaged. Only the central stripe of the primordium was imaged in S2E petal. For each, multiple Z-stacks (2.5 μm spacing) were acquired to cover the whole surface. For each petal, images stacks were stitched using ImageJ (Version 1.53q). Images were then analysed using MorphographX ^23^. The petal surface was extracted, and cell segmentation was performed, using the auto-segmentation function. Segmentation errors were corrected manually, and final meshes were converted into 2D meshes. The final template was processed with R and Matlab (Version 2020a) to extract the cell geometry information. To analyse the cell geometry distribution along the proximo-distal (PD) axis of the petal, a stripe of cells (20 % of the width of the petal at its widest point) in the central region of the petal was analysed.

#### Distribution of the cell features across the petal at S5

Images were acquired using a Keyence VHX-7000 digital microscope at 300X magnification. Cell features were manually measured along a line of cells, across the PD axis of the petal, using ImageJ (Version 1.53q) and processed using Matlab (Version 2020a).

#### Distribution of the cell division events across petals using EdU staining

For combined 5-ethynyl-2-deoxyuridine (Invitrogen A10044, Thermo Fisher Scientific) and modified pseudo-Schiff-propidium iodide (PI; Sigma-Aldrich Company) staining, the published method ^24^ has been modified. Buds from stage 0a to stage 1 were harvested and their sepals removed before being embedded in the EdU staining medium [0.22 % (w/v) Murashige and Skoog basal salts mixture, 3.5% (w/v) sucrose, 0.004% (w/v) L-cysteine, 0.0015% (w/v) ascorbic acid + 0.01% myo-inositol (w/v) + 0.0001% Nicotinic Acid (w/v) + 0.0001% (w/v) pyridoxine hydrochloride + 0.01% (w/v) thiamine hydrochloride + 0.0002% (w/v) glycine + 175 nm N6-Benzyladenine + 10 μM EdU (Invitrogen A10044, Thermo Fisher Scientific) pH 5.7] containing 0.8% (w/v) agarose. Liquid EdU staining medium was then added to immerse the bud. The samples were cultured 13h in a growth chamber (16 h light: 8 h dark photoperiod, average light intensity of 85 μM and average temperature of 24 °C). Petals were the dissected from the buds and all petals were dehydrated by successive 15 min treatments in an ethanol dilution series (15%, 30%, 50%, 70%, 85%, 95%, and 100% EtOH) and stored in 100% EtOH overnight. Samples were rehydrated through the same EtOH series and incubated at 37 °C overnight in alpha-amylase (Sigma-Aldrich A4551), 0.3 mg.mL^-1^ phosphate buffer 20 mM pH 7.0, 2 mM NaCl, 0.25 mM CaCl_2_. Petals were washed twice in water and once with Tris buffered saline (TBS) pH 7.4, before being incubated for 1h in solution containing 10 μM Alexa 488-azide (Invitrogen A10266, Thermo Fisher Scientific) and 100 mM Tris, pH 8.5; this was followed by 30 min in solution containing 10 μM Alexa 488-azide, 100 mM Tris, 1 mM CuSO4, 100 mM ascorbic acid, pH 8.5. Incubations were carried out at room temperature with gentle shaking and covered from the light. The samples were washed three times with water, treated in 1% periodic acid for 30 min, and washed twice with water, before being incubated in 0.01 μg.μL^-1^ propidium iodide for 3 hours, covered with gentle shaking. The petals were cleared with a chloral hydrate solution for 2 hours and mounted in Hoyer’s solution (30 g gum arabic, 200 g chloral hydrate, 20 g glycerol, and 50 mL water). Samples were imaged with a Zeiss LSM880 imaging system with 20x objective lens. Excitation at 488 nm and 561 nm; emission filters set to 499-526 nm for EdU and 603-629 nm for propidium iodide. For each sample from stage 0a to stage 1, multiple 3D Z-stacks were acquired to cover the whole petal. For stage 2, only a central stripe was imaged. Images were then processed using Imaris 9.2.1 (Bitplane). A stripe of 100 μm was selected in the centre of the petal and EdU-labeled nuclei were identified using the spot detection function (spot diameter: 4.16-5 μm). The data were then exported and analysed using Matlab (Version 2020a).

### Computational modelling

#### Data analysis

##### Boundary calculation

For the automatic calculation of the boundary between the two regions (used for the early pattern stages before the appearance of the pigmentation) from the data coming from MorphoGraphX in Fig. 5, we used the area as that is one of the defining characteristics for each cell type. For each sample, we first binned the cells in 5% windows based on their normalised position along the proximo-distal axis, *y*/ *L*, where *L* is the length of the tissue, to calculate an averaged cell-area along the proximo-distal axis. This window-averaged cell area is then further passed through a Savitzky–Golay filter to smooth the signal. Boundary position is then defined as the position of the average of all the windows within 2.5% of the window with the highest cell area.

##### Genotype comparison

To get the distribution of the ratios of the areas, number of cells, and boundaries we use the following definitions of the mean and variance of the ratio of two random variables, *X*and *Y*:

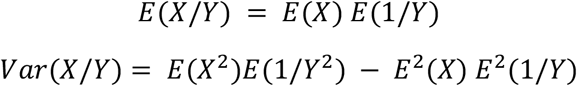

The error bars on the plots are the sample standard deviations,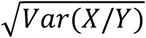.

##### Model description

Since the main question in this part of the work is the positioning of the boundary between cell types along the PD axis and our data are also mainly given in terms of their positioning along this axis, we represented the petal as a 1-D array of cells. This is a simplification, and it has been done for other tissues where one dimension of the tissue dominates, for example when considering hormone distributions in the root ^25^. The cells then grow and divide in a way that depends on their type and current size.

We used the modelling language Chromar which has been used before in cell-based simulations ^41^. Chromar uses discrete objects that carry attributes to represent entities in the model and rules on these objects to describe the dynamics. Rules are stochastic and the simulation is done with a version of the stochastic simulation algorithm (Gillespie – we refer the reader to the paper describing the language for details). In this case we used an implementation of this framework in Python (see Supplementary information for link to code – https://gitlab.developers.cam.ac.uk/slcu/teamem/lucie/tissue_hib_petal). Rules can also use aggregates over the state of the system called observables. Observables are made from two parts, a ‘selection’ part where one specifies which objects to choose from the state (e.g., all cells that have fate 0), and an aggregation part that defines how to reduce these cells into a single value (e.g., sum the length of the chosen cells).

In this case our objects are cells that have the following type:

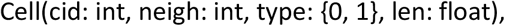

where cid is an integer cell identifier, neigh is an integer cell identifier representing the identity of their right neighbour, type is identifier for their type (fate), which in this case we choose to be either proximal or distal, and finally len is a float number representing their length. For notational convenience we use 0 for the type ‘distal’ and 1 for type ‘proximal’.

For *growth* we have the following rule:

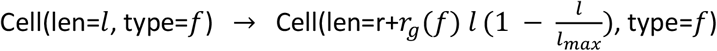

so, any cell in the state can grow its length by some amount that depends on its current length, its type (*r*_*g*_(*f*)) and how far away this length is from a maximal length that the cell can take (*l*_*max*_). The last two are functions of its type ^42^.

For *division* we have the following rule:

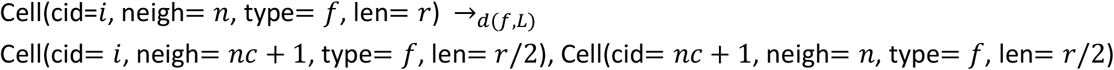

where,

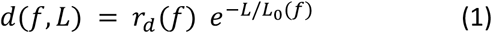

so, any cell can divide at a rate *d*(*f, L*) that depends on its fate and the current length of the whole petal. This rate depends on a basal division rate that depends on the fate of the cells, *r*_*d*_ (*f*), and a factor *e*^−*L*/*Lo* (*f*)^ that depends on the current length of the entire petal and a function of its type, *L*_*o*_(*f*), that controls the timing of a slow-down in the division rates that is fate-dependent. This slow-down of divisions over time is used to create the expansion phase of the development of the petal as observed in the data. Since the length of the petal is not in the attributes of the cell on the rule’s left-hand side, it needs to be computed with an *observable*:

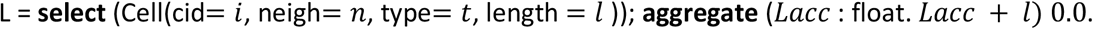

A division gives rise to a new cell to the right of the newly divided cell. In order to be able to give fresh ids to the new cells we need to know how many cells there are in total. So, we finally have the following *observable* to keep track of the number of cells in the tissue so we can give fresh ids to newly created cells:

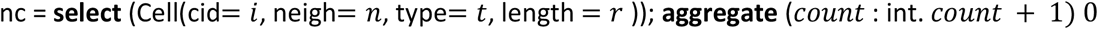

##### Model analysis

##### Initial and final state

For convenience for the initial state we started with 21 cells of length 0.1μm giving an initial length of around 2μm. From these cells the first 7 have the proximal fate and the rest have the distal fate. So, this assumes that even though there is no visible difference in the cell morphology, the cells have already assumed a fate in the beginning of the simulation and that the position of the fates along the PD axis is at the 1/3^rd^ position.

In the data we observe an increase in total length from 200 μm in stage S0a to about 30000μm in S5. For the model then we use a similar increase of ∼ x150 for a target total length of the petal of 300μm (Parameters, Table 1). The scaling down of the petal in terms of the number of cells in the simulation is done for computational efficiency reasons while not affecting the conclusions of the model.

**Table 1:** Estimation of the stage durations (approximation for stages 0a - 0b - 0c)

| Stage | Time (hr) |
| --- | --- |
| S0a | 1 |
| S0b | 31 |
| S0c | 62 |
| S1 | 93 |
| S2E | 152 |
| S2L | 211 |
| S3 | 309 |
| S4 | 404 |
| S5 | 416 |

**Table 2:** Main parameter values for the simulation in Fig. 6.

| Parameter | Values | Description |
| --- | --- | --- |
| $r_g(0)$ | 0.08 | base growth rate of distal cells, the base growth rate of proximal cells is given in reference to this |
| $r_d(0)$ | 0.06 | base division rate of distal cells, the base division rate of proximal cells is given in reference to this |
| $l_{max}$ | 20 | maximum length of cells |
| $L_0(0)$ | $0.11 * L_{max}$ | half-point for the decline in division rates as a function of the 'target' length of the petal. For the proximal fate this is given in terms of the distal $L_0$. |
| $L_{max}$ | 300 | threshold total length of petal |
| $L_0(0) / L_0(1)$ | 0.6 | difference in timing of exit from division-heavy phase for the two fates |

##### Parameters

Plotting the average growth dynamics of cell numbers and sizes (Fig. 5) allowed us to get a quantitative appreciation of the dynamics. To analyse the effect of model parameter values we first performed a more careful estimation of the growth parameters to fit with the data. For the growth and division parameters, (*r*_g_,*r*_d_), we estimate the rate for the distal cells from the data and then define the rate for the proximal cells relative to this value. The number of cells can be approximated to increase exponentially between states (Fig. 3f), giving an equation for the number of distal cells *dN*_0_ /*dt* =*kN*_0_, with solution 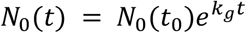, where *N*_*o*_(*t*_0_) is the initial number of distal cells and *N*_0_(*t*) is the number of distal cells at time *t*. The rate at time *t*_*i*_(a specific stage) is calculated as the rate of growth from the previous time *t*_*i*−1_(the previous stage) as 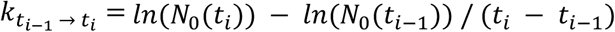. The timings of stages used in this calculation are given in Table 1 and the number of cells for each stage is the median of the cell numbers for that stage across the samples. No rate calculation was made for the first stage, S0a. Plotting the estimated rates against the length of the petal gives us an estimation of the parameters in *d*(0, *L*) (Eq. 1, Supplementary Fig. 7a), which are approximated using an exponential function.

Cell growth rates are harder to estimate since they are a combination of both an inherent ‘real’ growth rate of the cells and the rate of division. We can nevertheless, assuming an exponential growth as above (Supplementary Fig. 7b), estimate the effective growth rate of cells as 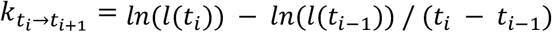, where *l*(*t*_*i*_) is the median cell area for all the cells for that stage across samples. The timings used for the stages are in Table 1 and as before there is no calculation for the first stage, S0a. The effective growth rate should be at its closest to the real growth rate towards the end of the process when the effect of divisions is at its lowest, we therefore used the calculated final effective growth rate as the true basal growth rate of the distal cells, *r*_*g*_(0).

##### Simulations

For each parameter choice referred to in the figures in the main text and elsewhere what changes is the ratio of the specific values of the basal growth rates per cell fate *r*_*g*_(0), *r*_*g*_(1) and *r*_*d*_(0), *r*_*d*_(1). All the simulations are run until the tissue reaches a certain threshold length. The total length of the tissue is the sum of the lengths of all the cells:

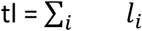

Since each simulation is stochastic for every parameter choice the model was run 100 times, then each of the interesting observables (number of cells in each region, growth rates in each region etc.) were binned into 1% of simulation-time windows to get 100 numbers per observable. These were then averaged across the 100 simulations.

##### Objective functions

To probe the behaviour of the model we used the dynamics of the position of the boundary, the area ratio – the ratio of average cell length in the two fates – and the number of cell ratio – the ratio of the number of cells in each fate as outputs. To calculate these from the state of the model, we define the following observables for the number of cells in each fate, nc0 and nc1, and the total length of each region, tl0 for the total length of the distal region and tl1 for the total length of the proximal region:

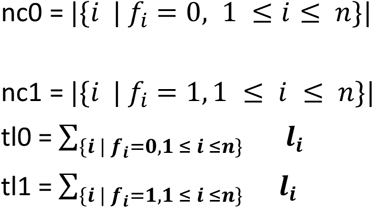

where *f*_*i*_ is the fate of cell *i, l*_*i*_is the length of cell and *n* is the total number of cells. The area ratio then is 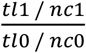, the number of cell ratio is 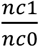 and the boundary position is 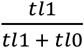These observables are calculated per step in the model output, binned into 1% of simulation-time windows and averaged to get 100 numbers. These were then averaged across 100 simulations. Since the length of the petal is a more accurate description of development and to ease the comparison to the experimental data, these were then resampled to every 10% of final length of the simulated petal to the nearest time-percentage. So, for example, the boundary position at 10% petal length (30, since the target length is 300) would be the value of the boundary position observable (calculated as above) at the time window where the length of the petal was nearest to that 10% of the total petal length.

In order to compare the model observables, the same observables had to be calculated for the experimental data. The calculated observables (Fig. 5) were interpolated and sampled to every 10% of final petal length as the model observables.

Finally, to compare a simulation to a data observable we used the mean percentage error. So, if the value of the model observable at 10% petal length is *obs*_*sim*_ and the value of the data observable at 10% petal length is *obs*_*data*_, the percentage error is (*obs*_*sim*_ − *obs*_*data*_) / *obs*_*data*_. These are then averaged across all samples. The total error for all three observables is the average of the three. This is what is plotted in Fig. 6c in the main text.

### Behavioural experiments

Flower-naïve bumblebees (*Bombus terrestris*) (Research Hive, UK) experiments were conducted using the same arena design as published ^19^. In total in this study, foragers from three colonies were used. Each colony was fed daily with fresh 15% sucrose solution and twice weekly with pollen (The Happy Health Company or Biobest). To distinguish individuals during experiments, foragers were hand-marked with water-based Thorne queen marking paint of various colour associations.

#### Artificial flower design

All artificial disks were 6 cm of diameter, made with a 3D-printer, using polylactic acid filament (1.75mm) and fixed, using Velcro® dots to a 3 cm cylinder of green FIMO® polyester clay. For the training phase, uniform black disks were used. For all differential conditioning, preference tests and foraging speed experiments, disks were bi-coloured with an outer white ring surrounding an inner purple circle of variable sizes depending on the desired bullseye dimensions: 1.2 cm (4%) in diameter for the small bullseye (*H. richardsonii*-like), 2.4 cm (16%) in diameter for the reference bullseye (medium size, *H. trionum*-like) and 3.6 cm (36%) in diameter for the larger bullseye (*35S::HtTCP4*.*1*-like).

#### Training phase

To familiarise themselves with the foraging set-up, bumblebees foraged freely from seven black artificial flowers (training discs), randomly positioned in the arena, each of them containing 45 μL of 15% w/w sucrose solution. After feeding, discs were removed, cleaned with 70% ethanol, and replaced in a new position. A bee was considered trained after it has made several return trips from the arena to the hive.

#### Differential conditioning experiments

During the test phase, two types of discs were compared at a time. Five artificial flower disks of one type (either small or large) plus five of the medium bullseye sizes were randomly positioned in the arena. Only one type of disc contained 45 μL of 20% sucrose solution (reward), the other type displayed 45 μL of ddH20 water (neutral reward). An individual trained bee was released into the arena, and that it visited were recorded. A disc was considered visited whenever a bee landed on it. After each visit, the disc was refilled with sucrose or water solution, and its position changed. Discs with which the bee came into contact were cleaned with 70% ethanol between each foraging bout and individual bees. For each bullseye size combination, experiments were performed with 20 bumblebees in total: e.g., for the comparison small vs. medium size, reward was presented on 10 bumblebees on the small disc and, reward was presented on the medium for 10 other ones. Each bee was tested up to a minimum of 80 choices. Statistical analyses were performed using RStudio (Version 1.1.1717). Learning curves associated with each pairwise comparison were obtained by pooling data from individual bees, as described in Moyroud et al., 2017.

#### Preference tests

Two types of preference tests were performed. The first one was a binary choice experiment, where a naïve bumblebee was presented with two discs at equidistant entrance of the hive, and equally rewarding (45 μL of 20% sucrose solution). The preference of 30 individuals were recorded for each type of bullseye size combination (small *vs*. medium, and medium *vs*. large). Among this, 15 bumblebees were presented the first type on the right, and the 15 others on the left. Statistical differences were calculated using a one sample t-test (RStudio (Version 1.1.1717)). The second preference test was subsequently performed to identify consistency in preference over 10 choices in a larger foraging display. Ten rewarding discs, five of each bullseye size, were presented simultaneously and the ten first choices made by an individual forager were recorded. Statistical analyses were performed using RStudio (Version 1.4.1717), using a two-sided t-test, assessing significant increase in % of one disc-type chosen from than would be expected at random (50%).

#### Foraging speed experiments

Three artificial flowers, all from the same type (bullseye size from type 1), and offering 30 μl of 30% sucrose solution, were set 30 cm apart from one another in the arena (in position 1). A naïve individual was introduced to the arena and the times it took to move between the three discs (disc 1 to disc 2, then disc 2 to disc 3) during a foraging bout were recorded with a Samsung Galaxy Tab E tablet. A large reward (100 μl of 30% sucrose solution) was offered to the bee at the end of the foraging bout to encourage it to return to the hive. Flowers with which the bees came into contact were cleaned with 70% ethanol, and then water to remove scent marks. Then, the same bumblebee was offered to repeat the same experiment with the second type of artificial flowers (bullseye size, type 2), set 30 cm apart from one another in a new location (position 2). The same experiment, with the same bee, was repeated with the third disc type (type 3) at a new position in the arena (position 3). The entire procedure was then repeated at least five times to ensure that 5 complete foraging bouts on each flower type were recorded for each individual bee (15 foraging bouts total for each bee). This routine allowed us to control the variability in foraging speed between foragers (as each bee performed the experiment on each type of flower) and any potential effect of the position of the flowers in the arena. In total, 15 individuals were independently tested.

The time taken for each bee to travel between each disc was extracted from the recordings using VLC Video Software and the Time v3.2 extension. After examining the plots of residuals, a single-factor ANOVA and a post hoc Tukey’s HSD test were conducted in RStudio (Version 1.4.1717) to explore differences between artificial flower types.

## Acknowledgments

We thank Prof. B. G. Murray for the generous gift of *H. richardsonii* seeds, Dr Katharina Schiessl for advice regarding the EdU staining protocol, Dr. Raymond Wightman and Mr. Gareth Evans for their assistance with confocal imaging and Dr. Elena Salvi for guidance and expertise with the IMARIS software. We also would like to thank all members of team Hibiscus at SLCU for their engagement with discussions and feedback on the results presented here and the entire professional service community at SLCU. We thank in particular the general lab support team for media preparation, the horticulture team for their help with plant care and pest control and the facilities team for their help setting up the bee lab and building our flight arenas.

## Author contributions

EM and LR designed the study; LR, ALF, MTY and EM performed the experiments as follows: LR established the confocal imaging pipeline, developed the EdU staining protocol and performed the image acquisition, processing and analyses. EM constructed the *35S::HtTCP4*.*1* plant expression plasmid and generated the corresponding *H. trionum* transgenic lines. MTY constructed the *35S::HtTCP4*.*2* plant expression plasmid and generated the corresponding transgenic lines. MTY characterised the bullseye phenotype of both *HtTCP4* transgenic lines, LR performed the cell feature analysis. LR, EM and ALF performed the bee behavioural experiments and analysed the data. AZ and HJ designed the computational model. AZ performed the model simulations and evolution of ratio quantifications. LR prepared the figures with the help of EM and AZ. LR and EM wrote the manuscript. All authors read and commented on successive drafts of the manuscript and approved the submitted version.

## References

1. Davies, K. M., Albert, N. W,. & Schwinn, K. E. From landing lights to mimicry: the molecular regulation of flower colouration and mechanisms for pigmentation patterning. Funct Plant Biol 39, 619–638 (2012).

2. Fairnie, A. L. M. et al. Eco-Evo-Devo of petal pigmentation patterning. Essays in Biochemistry 66, 753–768 (2022).

3. Rausher, M. D. Evolutionary Transitions in Floral Color. International Journal of Plant Sciences 169, 7–21 (2008).

4. Gao, S. et al. Effects of drought stress on growth, physiology and secondary metabolites of Two Adonis species in Northeast China. Scientia Horticulturae 259, 108795 (2020).

5. Li, B. et al. Flavonoids improve drought tolerance of maize seedlings by regulating the homeostasis of reactive oxygen species. Plant Soil 461, 389–405 (2021).

6. Rao, M. J. et al. CsCYT75B1, a Citrus CYTOCHROME P450 Gene, Is Involved in Accumulation of Antioxidant Flavonoids and Induces Drought Tolerance in Transgenic Arabidopsis. Antioxidants 9, 161 (2020).

7. Talbi, S. et al. Effect of drought on growth, photosynthesis and total antioxidant capacity of the saharan plant Oudeneya africana. Environmental and Experimental Botany 176, 104099 (2020).

8. Todesco, M. et al. Genetic basis and dual adaptive role of floral pigmentation in sunflowers. eLife 11, e72072 (2021).

9. Koski, M. H,. & Ashman, T.-L. Floral pigmentation patterns provide an example of Gloger’s rule in plants. Nature Plants 1, 1–5 (2015).

10. Koski, M. H., MacQueen, D,. & Ashman, T.-L. Floral Pigmentation Has Responded Rapidly to Global Change in Ozone and Temperature. Curr Biol 30, 4425-4431.e3 (2020).

11. Ding, B. et al. Two MYB Proteins in a Self-Organizing Activator-Inhibitor System Produce Spotted Pigmentation Patterns. Current Biology 30, 802-814.e8 (2020).

12. Lin, R.-C,. & Rausher, M. D. R2R3-MYB genes control petal pigmentation patterning in Clarkia gracilis ssp. sonomensis (Onagraceae). New Phytologist 229, 1147–1162 (2021).

13. Martins, T. R., Berg, J. J., Blinka, S., Rausher, M. D,. & Baum, D. A. Precise spatio-temporal regulation of the anthocyanin biosynthetic pathway leads to petal spot formation in Clarkia gracilis (Onagraceae). New Phytologist 197, 958–969 (2013).

14. Sagawa, J. M. et al. An R2R3-MYB transcription factor regulates carotenoid pigmentation in Mimulus lewisii flowers. New Phytologist 209, 1049–1057 (2016).

15. Shang, Y. et al. The molecular basis for venation patterning of pigmentation and its effect on pollinator attraction in flowers of Antirrhinum. New Phytologist 189, 602–615 (2011).

16. Yuan, Y.-W., Rebocho, A. B., Sagawa, J. M., Stanley, L. E,. & Bradshaw, H. D. Competition between anthocyanin and flavonol biosynthesis produces spatial pattern variation of floral pigments between Mimulus species. Proceedings of the National Academy of Sciences 113, 2448–2453 (2016).

17. Riglet, L., Gatti, S,. & Moyroud, E. Sculpting the surface: Structural patterning of plant epidermis. iScience 24, 103346 (2021).

18. Moyroud, E. et al. Cuticle chemistry drives the development of diffraction gratings on the surface of Hibiscus trionum petals. Current Biology 32, 5323-5334.e6 (2022).

19. Moyroud, E. et al. Disorder in convergent floral nanostructures enhances signalling to bees. Nature 550, 469–474 (2017).

20. Vignolini, S. et al. The flower of Hibiscus trionum is both visibly and measurably iridescent. New Phytologist 205, 97–101 (2015).

21. Whitney, H. M. et al. Floral Iridescence, Produced by Diffractive Optics, Acts As a Cue for Animal Pollinators. Science 323, 130–133 (2009).

22. Craven, L., de Lange, P., Lally, T., Murray, B,. & Johnson, S. A taxonomic re-evaluation of Hibiscus trionum (Malvaceae) in Australasia. New Zealand Journal of Botany 49, 27–40 (2011).

23. Barbier de Reuille, P. et al. MorphoGraphX: A platform for quantifying morphogenesis in 4D. eLife 4, e05864 (2015).

24. Schiessl, K. et al. NODULE INCEPTION Recruits the Lateral Root Developmental Program for Symbiotic Nodule Organogenesis in Medicago truncatula. Current Biology 29, 3657–3668.e5 (2019).

25. Mironova, V. V. et al. A plausible mechanism for auxin patterning along the developing root. BMC Systems Biology 4, 98 (2010).

26. Cubas, P., Lauter, N., Doebley, J,. & Coen, E. The TCP domain: a motif found in proteins regulating plant growth and development. Plant J 18, 215–222 (1999).

27. Doebley, J., Stec, A,. & Hubbard, L. The evolution of apical dominance in maize. Nature 386, 485–488 (1997).

28. Kosugi, S,. & Ohashi, Y. PCF1 and PCF2 specifically bind to cis elements in the rice proliferating cell nuclear antigen gene. Plant Cell 9, 1607–1619 (1997).

29. Luo, D., Carpenter, R., Vincent, C., Copsey, L,. & Coen, E. Origin of floral asymmetry in Antirrhinum. Nature 383, 794–799 (1996).

30. Goulson, D,. & Hanley, M. E. Distribution and forage use of exotic bumblebees in South Island, New Zealand. New Zealand Journal of Ecology 28, 225–232 (2004).

31. Schmid-Hempel, P., Schmid-Hempel, R., Brunner, P. C., Seeman, O. D,. & Allen, G. R. Invasion success of the bumblebee, Bombus terrestris, despite a drastic genetic bottleneck. Heredity 99, 414–422 (2007).

32. Galipot, P., Damerval, C,. & Jabbour, F. The seven ways eukaryotes produce repeated colour motifs on external tissues. Biological Reviews 96, 1676–1693 (2021).

33. Burian, A. et al. Specification of leaf dorsiventrality via a prepatterned binary readout of a uniform auxin input. Nat. Plants 8, 269–280 (2022).

34. Caggiano, M. P. et al. Cell type boundaries organize plant development. eLife 6, e27421 (2017).

35. Lewis, M. W,. & Hake, S. Keep on growing: building and patterning leaves in the grasses. Current Opinion in Plant Biology 29, 80–86 (2016).

36. Liu, J., Moore, S., Chen, C,. & Lindsey, K. Crosstalk Complexities between Auxin, Cytokinin, and Ethylene in Arabidopsis Root Development: From Experiments to Systems Modeling, and Back Again. Mol Plant 10, 1480–1496 (2017).

37. Müller, C. J., Larsson, E., Spíchal, L,. & Sundberg, E. Cytokinin-Auxin Crosstalk in the Gynoecial Primordium Ensures Correct Domain Patterning. Plant Physiol 175, 1144–1157 (2017).

38. Challa, K. R., Aggarwal, P,. & Nath, U. Activation of YUCCA5 by the Transcription Factor TCP4 Integrates Developmental and Environmental Signals to Promote Hypocotyl Elongation in Arabidopsis. Plant Cell 28, 2117–2130 (2016).

39. Murray, B. G., Craven, L. A,. & De Lange, P. J. New observations on chromosome number variation in Hibiscus trionum s.l. (Malvaceae) and their implications for systematics and conservation. New Zealand Journal of Botany 46, 315–319 (2008).

40. Gibson, D. G. et al. Enzymatic assembly of DNA molecules up to several hundred kilobases. Nat Methods 6, 343–345 (2009).

41. Honorato-Zimmer, R., Millar, A. J., Plotkin, G. D,. & Zardilis, A. Chromar, a language of parameterised agents. Theoretical Computer Science 765, 97–119 (2019).

42. Jönsson, H., Heisler, M. G., Shapiro, B. E., Meyerowitz, E. M,. & Mjolsness, E. An auxin-driven polarized transport model for phyllotaxis. Proceedings of the National Academy of Sciences 103, 1633–1638 (2006).

