## Supplementary Figures 1 to 7 for "Hibiscus bullseyes reveal mechanisms controlling petal pattern proportions that influence plant-pollinator interactions"

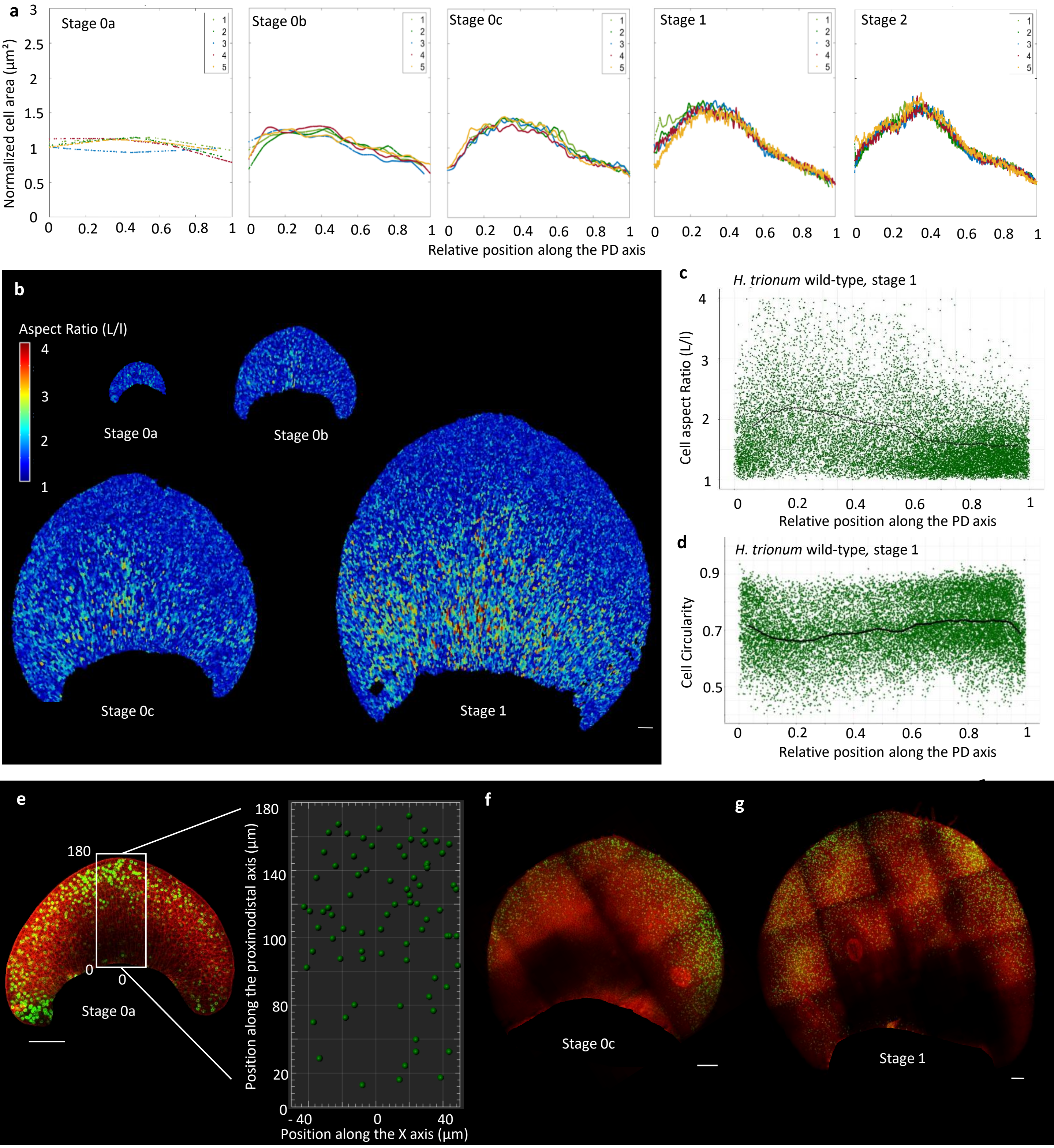


**Supplementary Fig. 1. Characterisation of cell behaviour across the adaxial petal epidermis of *H. trionum* wild-type during early development**

**a**, Normalised cell area distribution (cell area/mean of the cell area) across the proximo-distal axis of wild-type *H. trionum* petal. Only the central stripe (20% of the petal width) was taken into account in these graphs to maximize readability. Cell positions along the PD axis are relative, with 0 corresponding to the petal base, and 1 to the petal tip. Each line corresponds to the normalised cell area for one replicate. n=5 petals for each stage. **b**, Colour map of the cell aspect ratio of *H. trionum* wild-type petal. At S1, the cells in the proximal region, which will become the elongated and flat purple cells at S5, already appear longer and thinner than the ones in the distal region. Scale bar, 100 µm. **c**, Cell aspect ratio (major L/minor l) across the PD axis of the *H. trionum* wild-type petal at S1. Each green dot represents a cell, n=5 petals. **d,** Cell circularity across the PD axis of the *H. trionum* wild-type petal at S1. Each green dot represents a cell. n=5 petals. **e**, Distribution of cell division events across the PD axis of the *H. trionum* wild-type petal at S0a, using EdU staining to label newly replicated DNA (green) and propidium iodide to label the plasma membrane (red). Quantification of the EdU-labelled nuclei using Imaris, on a central stripe (white rectangle), along the proximo-distal axis of the petal. Example for one S0a petal. Division events occur mostly in the distal region of the petal. Scale bar, 100 µm **f**, As in (e) but for S0c petals of wild-type *H. trionum*. Scale bar, 100 µm **g**, As in (e), but for S1 petals of wild-type *H.trionum*. Scale bar, 100 µm a

b

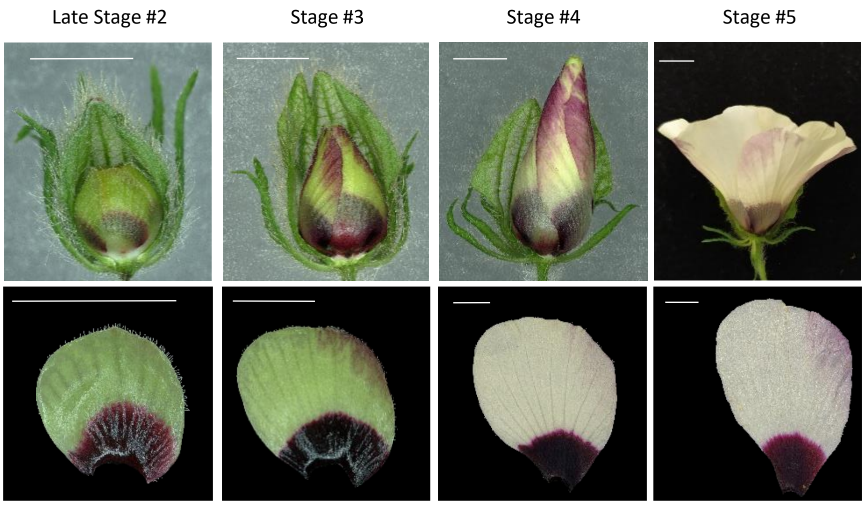


c

| | Late Stage #2 | Stage #3 | Stage #4 | Stage #5 |
|---|---|---|---|---|
| Range | 3000 - 5500 µm | 5500 – 12000 µm | 12000 – 23000 µm | 28000 - 33000 µm |
| Petal length (µm) | 4008 (+/-409) | 7144 (+/-1074) | 18096 (+/-2152) | 30564 (+/-1886) |
| Total cell number on a stripe of cells | 416 (+/-70) | 701 (+/-47) | 784 (+/-47) | 827 (+/-65) |

d

e

f

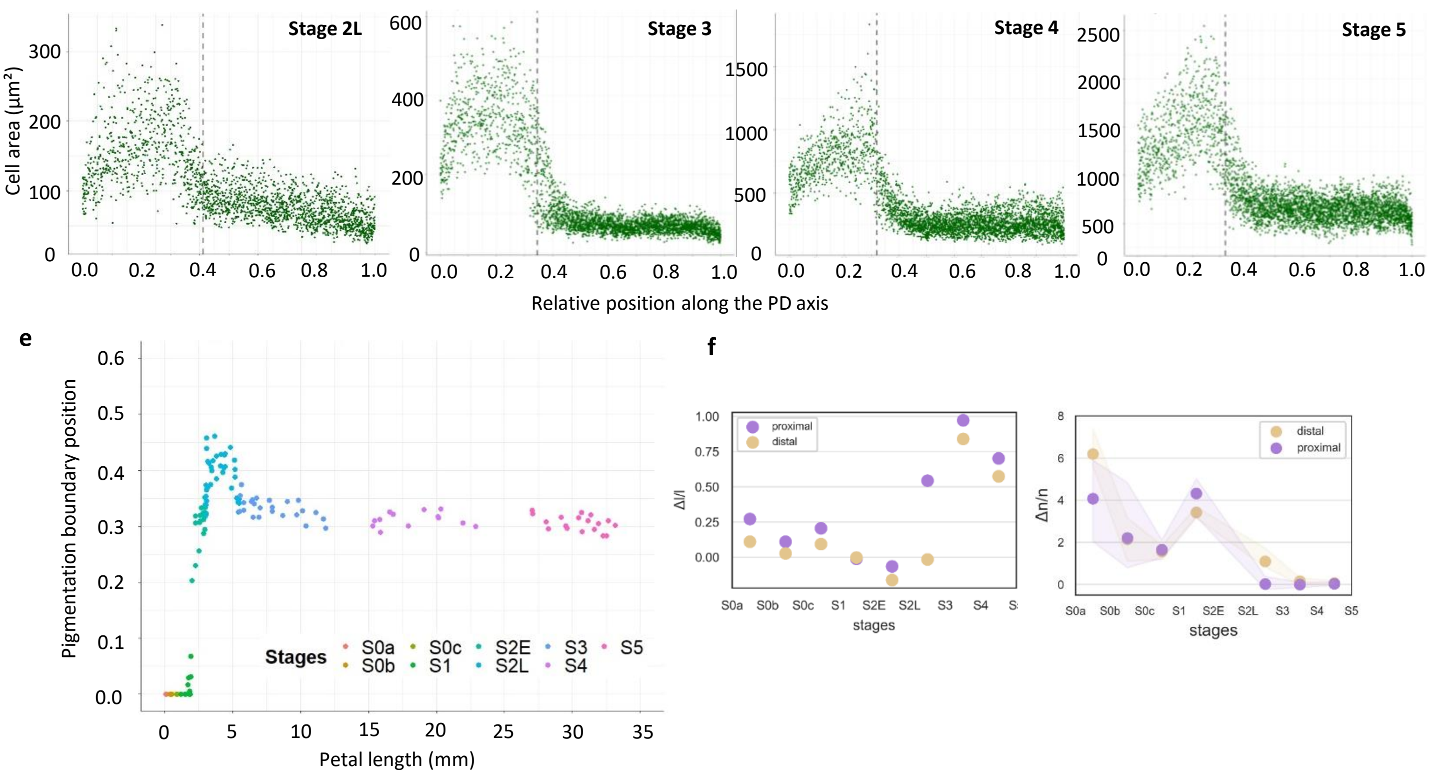


**Supplementary Fig. 2. Petal dimensions, bullseye boundary position and cell area distribution across the adaxial petal epidermis during later developmental stages (Late S2 to S5) of wild-type *Hibiscus trionum* .**

**a**, Petals on the *H. trionum* flower buds from Late Stage 2 to Stage 5, half of the sepals were removed to expose the petals. Only the abaxial side of each petal is visible here. Scale bars, 5 mm. **b**, View of the adaxial petal epidermis during later developmental stages (Late Stage 2 to Stage 5). Scale bars, 5 mm. **c**, Classification criteria for staging of *Hibiscus trionum* petal primordia during later developmental phase (from Late Stage 2 to Stage 5). The total number of cells was counted on a central stripe representing 20% of the petal width. **d**, Cell area distribution across the proximo-distal axis of *H. trionum* petal from Late Stage 2 to Stage 5. Only a central line of cells was measured. Cell positions along the PD axis are relative, with 0 corresponding to the petal base, and 1 to the petal tip. The dashed line corresponds to the average position of the pigmentation boundary for each stage. **e**, Relative position (0 corresponding to the petal base, and 1 to the petal tip) of the bullseye boundary (pigmentation transition) throughout petal development in *H. trionum* wild-type. n=5 petals for all stages.**f**, Rates of median (left) cell expansion and (right) cell division per petal domains in *H. trionum* WT throughout petal morphogenesis.

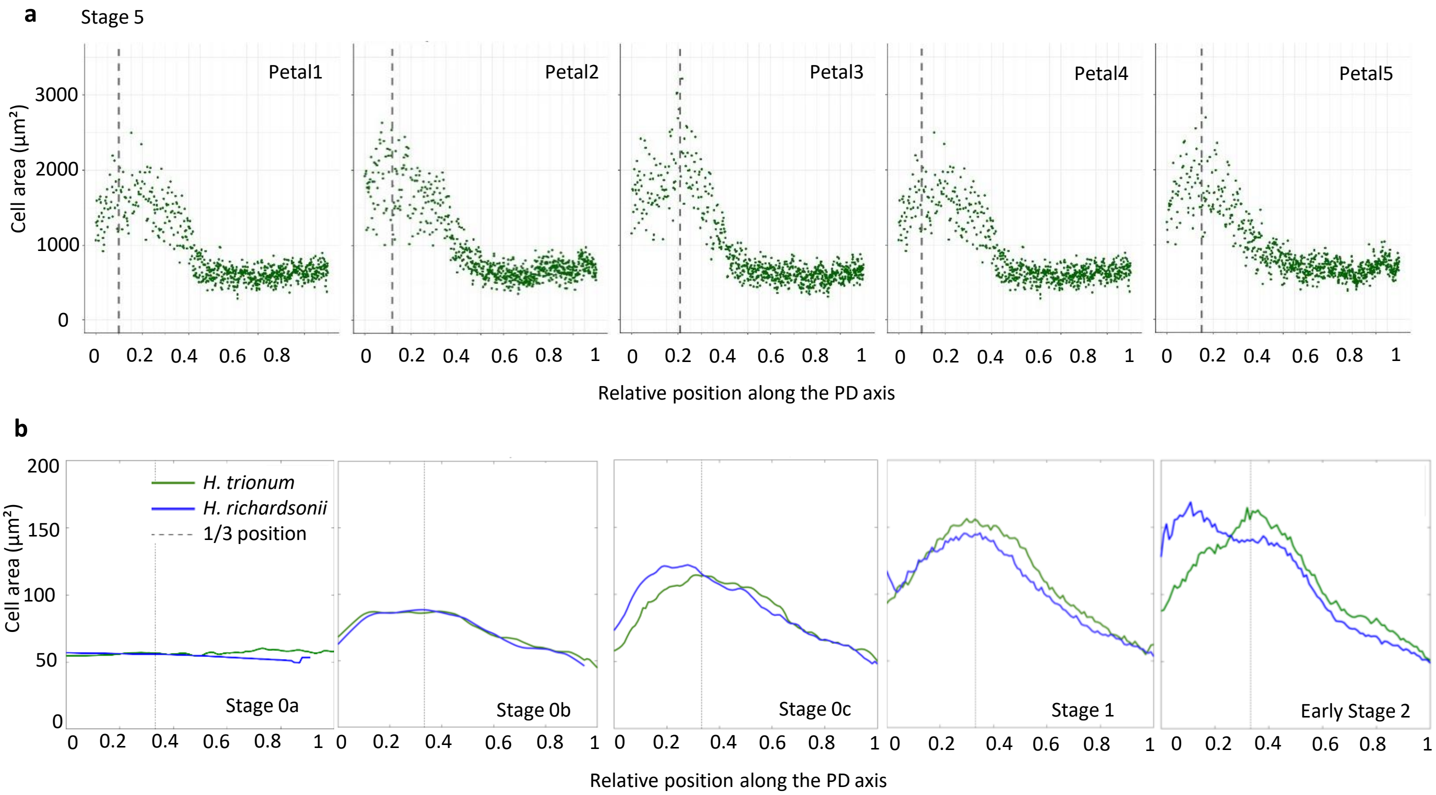


**Supplementary Fig. 3. Distribution of cell area across the adaxial petal epidermis of *H. richardsonii* during petal development.**
**a**, Distribution of the cell area across the PD axis of the *H. richardsonii* petal at S5 (mature flower). Each green dot represents a cell, each graph represents a replicate (n=5 petals). The dashed line corresponds to the peak of larger cells position. **b**, Cell area distribution across the proximo-distal axis of *H. richardsonii* (blue) and *H. trionum* wild-type (green) petals. Only the central stripe (20% of the petal width) was taken into account in these graphs to maximize readability. Cell positions along the PD axis are relative, with 0 corresponding to the petal base, and 1 to the petal tip. Each line corresponds to the average cell area of all replicates for each stage in each species. n=5 petals for each stage.

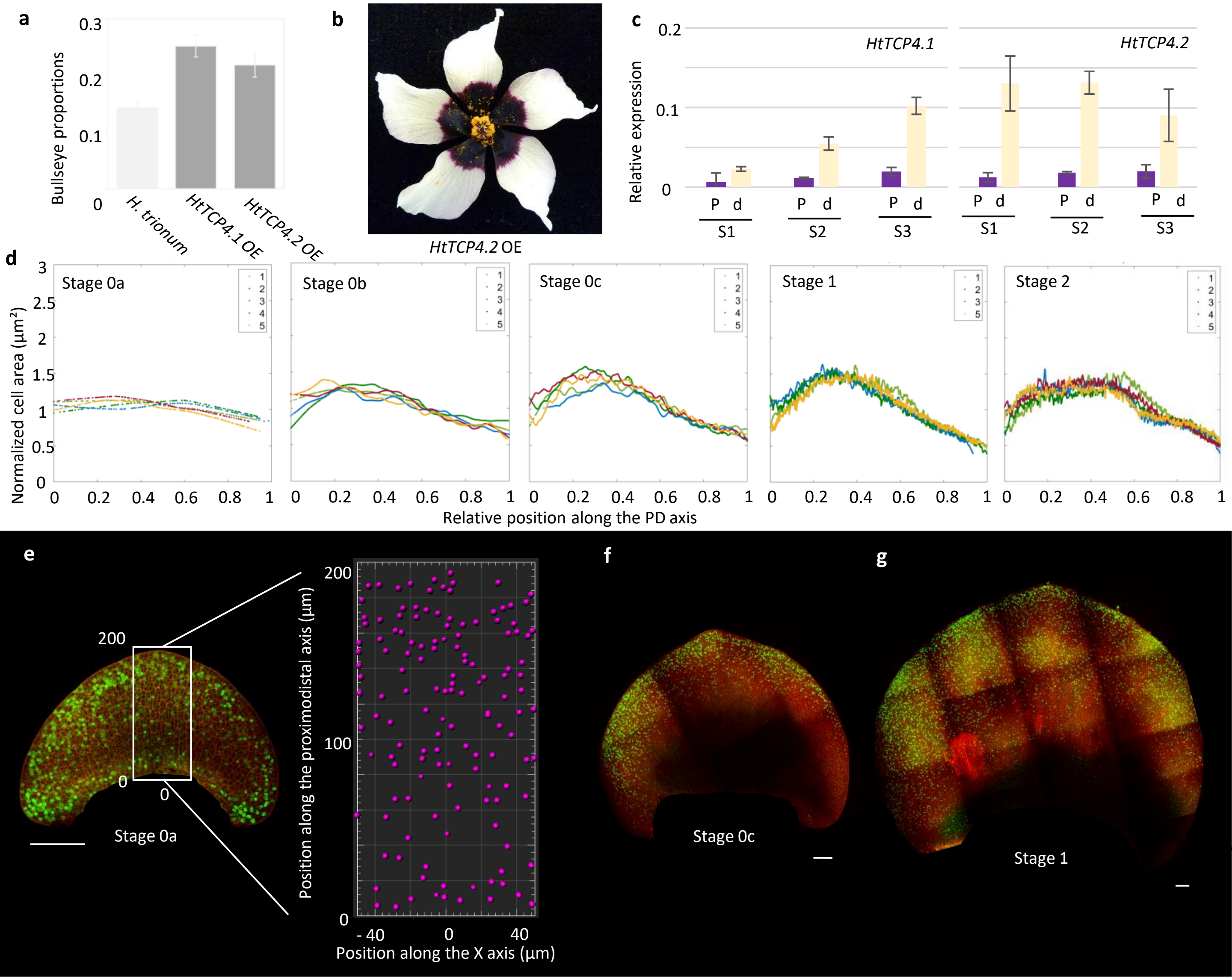


**Supplementary Fig. 4. Bullseye proportions in transgenic lines overexpressing *HtTCP4.1* or *HtTCP4.2* and characterisation of cell behaviour across the adaxial petal epidermis of *H. trionum HtTCP4.1* OE transgenic line during early development.**

**a**, Bullseye proportions in *H. trionum* wild-type, transgenic line *HtTCP4.1* OE and transgenic line *HtTCP4.2* OE . **b**, Opened flower of transgenic line *HtTCP4.2* OE. **c**, Relative expression of *HtTCP4.1* and *HtTCP4.2* in *H. trionum* wild-type from S1 to S4, in the proximal (P) and distal (d) regions of the petal. **d**, Normalised cell area distribution (cell area/mean of the cell area) across the proximo-distal axis of *HtTCP4.1* OE *H. trionum* petal. Only the central stripe (20% of the petal width) was taken into account in these graphs to maximize readability. Cell positions along the PD axis are relative, with 0 corresponding to the petal base, and 1 to the petal tip. Each line corresponds to the normalised cell area for one replicate. n=5 petals for each stage (except for S1 for which n=4). **e**, Distribution of cell division events across the PD axis of the *H. trionum HtTCP4.1* OE petal at S0a, using EdU staining to label newly replicated DNA (green) and propidium iodide to label the plasma membrane (red). Quantification of the EdU-labelled nuclei using Imaris, on a central stripe (white rectangle), along the proximo-distal axis of the petal. Example for one S0a petal. Division events occur across the entire petal adaxial epidermis. Scale bar, 100 µm. **f**, As in (e) but for S0c petals of *H. trionum HtTCP4.1* OE. Scale bar, 100 µm. **g**, As in (e), but for S1 petals of *H.trionum HtTCP4.1* OE. Scale bar, 100 µm.

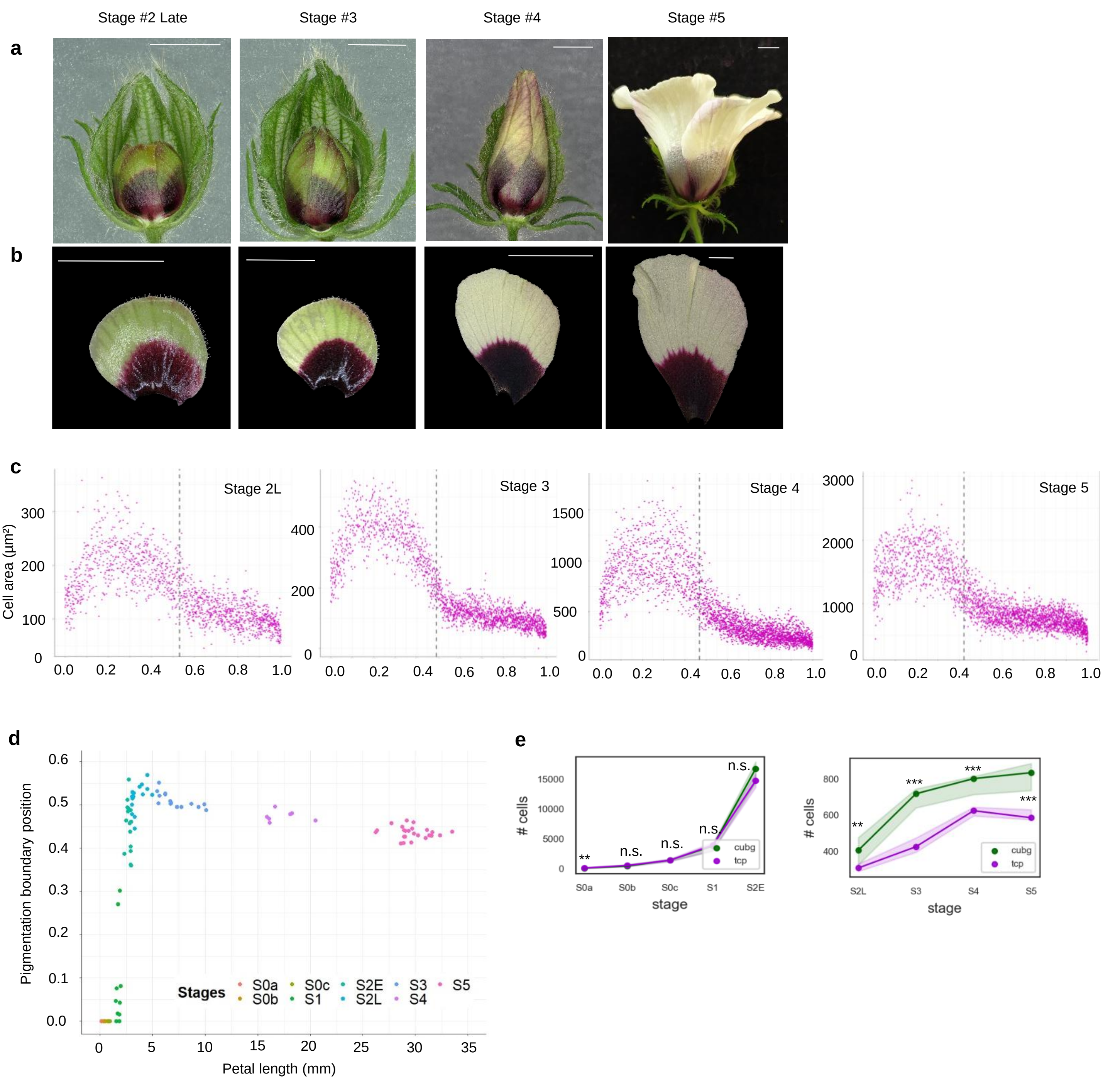


**Supplementary Fig. 5.** ***35S::HtTCP4.1*** **later developmental stages**

**a**, Petals on the *H. trionum HtTCP4.1* OE transgenic line flower buds from Late Stage 2 to Stage 5, half of the sepals were removed to expose the petals. Only the abaxial side of each petal is visible here. Scale bars, 5 mm. **b**, View of the adaxial petal epidermis in *H. trionum HtTCP4.1* OE transgenic line during later developmental stages (Late Stage 2 to Stage 5). Scale bars, 5 mm. **c**, Cell area distribution across the proximo-distal axis of *H. trionum HtTCP4.1* OE transgenic line petal from Late Stage 2 to Stage 5. Only a central line of cells was measured. Cell positions along the PD axis are relative, with 0 corresponding to the petal base, and 1 to the petal tip. The dashed line corresponds to the average position of the pigmentation boundary for each stage. n=5 petals for each stage. **d**, Relative position (0 corresponding to the petal base, and 1 to the petal tip) of the bullseye boundary (pigmentation transition) throughout petal development in *H. trionum HtTCP4.1* OE transgenic line. n=5 petals for all stages. **e**, Comparison of cell numbers across the adaxial petal epidermis of (green) *H. trionum* wild-type and (purple) *HtTCP4.1* OE transgenic line throughout the early phase (S0a to Early S2) or late phase (Late S2 to S5) of petal development. For the late phase cell numbers were only computed along a single line of cells, central to the petal. **$p<0.05$, ***$p<0.01$, n.s. = non significant.

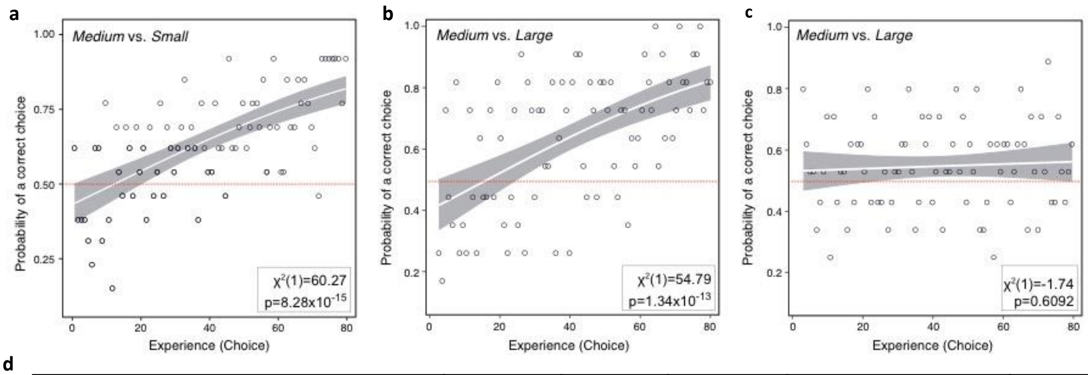


**d**

| | | | t | df | p-value | 95% confidence interval | Mean | n |
|---|---|---|---|---|---|---|---|---|
| ***BINARY TESTS*** | Medium vs. Small | | 3.395 | 29 | 0.002005 | 0.606- 0.927 | 0.76 | 30 bees |
| | Medium vs. Large | | 1.099 | 29 | 0.281 | 0.414-0.786 | 0.6 | 30 bees |
| ***10 CHOICE TESTS*** | Medium vs. Small | First five choices | 6.243 | 14 | 2.151e-05 | 3.266 - 4.067 | 3.7 | 15 bees |
| | | First ten choices | 7.897 | 14 | 1.592e-06 | 6.360 - 7.373 | 6.9 | 15 bees |
| | Medium vs. Large | First five choices | 0.468 | 14 | 0.647 | 2.141 - 3.059 | 2.6 | 15 bees |
| | | First ten choices | 0.899 | 14 | 0.384 | 4.723- 5.677 | 5.2 | 15 bees |

**Supplementary Fig. 6 Impact of bullseye size on bumblebees (*Bombus terrestris*) behaviour**

**a**, Differential conditioning experiment – comparison between Medium vs. Small bullseye size. In this graph, only 13 bumblebees out of the 20 from Fig. 7b were included: 5 bees were removed since they had already a strong preference for the medium-size bullseye (wild-type *H. trionum*-like) from the start of the experiment, and 2 did not appear to learn. **b**, As Fig. 7c but only including 11 bumblebees (out of the 22 tested) that learnt to distinguish between the discs with Medium (wild-type *H. trionum*-like) vs. Large bullseyes (*H. trionum HtTCP4.1* OE-*like*). **c**, As in **b**, but with the remaining 11 bees from Fig. 7c that fail to show any ability to distinguish between Medium and Large bullseyes. (**d**) Statistics table from the preference tests experiments. *First row:* Binary test. bumblebees showed a statistically significant preference for the medium (wild-type *H. trionum*-like) bullseye size compared to the small one (*H. richardsonii*-like), and no significant difference for the medium *vs.* the larger bullseye (one sample t-test). *Second row:* 10 choice tests. When only the first five choices or all 10 choices were considered, bumblebees showed a statistically significant preference for the medium (wild-type *H. trionum*-like) bullseye size compared to the small one (*H. richardsonii*-like), and no significant difference for the medium (wild-type *H. trionum*-like) *vs.* the large bullseye (*H. trionum HtTCP4.1* OE-*like*).

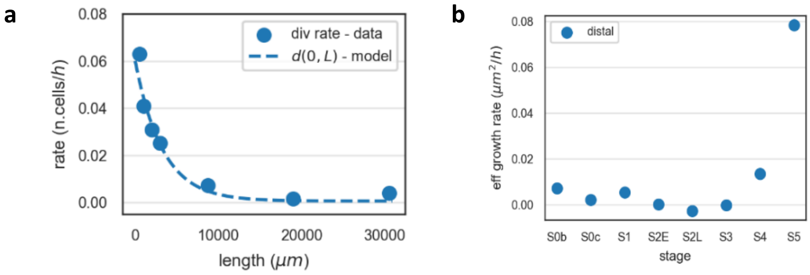


**Supplementary Fig. 7. Parameter estimations for the computational simulations. a,** Estimation of division rate in the distal region in wild-type *H. trionum*. These estimates informed the parameters choice in Table 1 (Materials and Methods section). **b,** Estimation of the effective growth rate of the distal region in wild-type *H. trionum.* These estimates informed the parameters choice in Table 1 (Materials and Methods section).
